# Voltage- and Branch-specific Climbing Fiber Responses in Purkinje Cells

**DOI:** 10.1101/284026

**Authors:** Yunliang Zang, Stéphane Dieudonné, Erik De Schutter

## Abstract

Climbing fibers (CFs) provide instructive signals driving cerebellar learning, but mechanisms causing the variable CF responses in Purkinje cells (PCs) are not fully understood. Using a new experimentally validated PC model, we unveil the ionic mechanisms underlying CF-evoked distinct spike waveforms on different parts of the PC. We demonstrate that voltage can gate both the amplitude and the spatial range of CF-evoked Ca^2+^ influx by the availability of K^+^ currents. This makes the energy consumed during a complex spike (CS) also voltage-dependent. PC dendrites exhibit inhomogeneous excitability with individual branches as computational units for CF input. The variability of somatic CSs can be explained by voltage state, CF activation phase, and instantaneous CF firing rate. Concurrent clustered synaptic inputs affect CSs by modulating dendritic responses in a spatially precise way. The voltage- and branch-specific CF responses can increase dendritic computational capacity and enable PCs to actively integrate CF signals.

## Introduction

According to the Marr-Albus-Ito theory (Albus, 1971; Ito, 1972; Marr, 1969), climbing fiber (CF) inputs to Purkinje cells (PCs) carry movement error information and evoke cerebellar learning by depressing the strength of parallel fiber (PF) synaptic input. Recently, processing of CF-carried error information by PCs has become a focus of cerebellar research (Najafi et al., 2014; Yang and Lisberger, 2014), with increased attention to modifiable dendritic responses (Davie et al., 2008; Kitamura and Hausser, 2011; Najafi et al., 2014; Rokni et al., 2009). Unfortunately, the biophysical mechanisms underlying the voltage-related dendritic response amplitude are still unresolved. CF provides powerful synaptic input onto proximal dendrites, but Ca^2+^ influx in distal dendrites is found to be unreliable (Ohtsuki et al., 2012; Otsu et al., 2014; Zagha et al., 2010). The pattern of dendritic spike initiation and propagation is important because the spatial range of Ca^2+^ influx controls the dendritic sites undergoing synaptic plasticity and even the direction of cerebellar learning. Moreover, whether individual branches of PC dendritic trees are homogeneously excitable has never been investigated due to technical limitations. The effect of clustered PF synaptic input (Wilms and Hausser, 2015) on CF-evoked dendritic responses is also unknown. The answers to these questions will determine whether the computational unit of CF responses is the entire dendritic tree or individual branches.

Somatic complex spikes (CSs) were assumed to be a result of synaptic input current and somatic ionic currents (Schmolesky et al., 2002). Observations that dendritic spike variation plays a minimal role in evoking an extra somatic spikelet seemed to confirm this assumption (Davie et al., 2008), however, contrary observations exist (Ohtsuki et al., 2012; Otsu et al., 2014). Somatic CSs are quite variable (Burroughs et al., 2017; Warnaar et al., 2015) and their durations have been linked with the degree of trial-over-trail learning (Yang and Lisberger, 2014). Voltage states or simple spike firing rates (SSFRs) (well known to represent voltage states *in vitro* and recently demonstrated *in vivo* (Jelitai et al., 2016)) have been correlated with CS spikelet numbers in some studies (Burroughs et al., 2017; Gilbert, 1976; Khaliq and Raman, 2005; Monsivais et al., 2005), but not in others (Mano, 1970; Warnaar et al., 2015). Additionally, CSs can be modulated by CF FRs (Hansel and Linden, 2000), but contradictory correlations between spikelet numbers and the instantaneous CF FRs have been reported (Burroughs et al., 2017; Khaliq and Raman, 2005; Maruta et al., 2007; Warnaar et al., 2015). Thus, it is necessary to re-examine how the CS shape is regulated.

Spiking is energetically expensive (Attwell and Laughlin, 2001). Each CS consumes more energy than a SS, but neither can be measured experimentally.

To address these questions, a systematic exploration of CF responses with high spatio-temporal resolution is required. Given the difficulty of simultaneously recording at numerous locations in the same cell by patch-clamp (Davie et al., 2008; Stuart and Hausser, 1994) and the lack of direct spike information from Ca^2+^ imaging (Deneux et al., 2016), computer models can play an indispensable role. Here we have built a ‘canonical’ PC model that reproduces most available experimental observations instead of fitting it to a specific cell recording. We use this model to systematically explore factors that affect CF-evoked dendritic and somatic responses and estimate the energy consumed for SSs and CSs.

## Results

### Electrophysiological properties of SSs

Before using the model to make predictions, we first compare its basic properties with *in vitro* experimental data. Example cell recordings are shown in Fig. 1A-D to help the comparison. Our PC model spontaneously fires at 40 Hz in the absence of synaptic inputs or current injections (27 to 140 Hz in experiments). SSs initiate first at the axon initial segment (AIS) with an axosomatic delay of 0.12 ms (~ 0.1 ms reported by Palmer et al. (2010)). The SS peak is -1 mV and its amplitude is 68 mV (−11 to −3 mV and 56 to 68 mV respectively in experiments). SSs have a half-amplitude duration of 0.22 ms (0.22 to 0.28 ms in experiments). The peak *dv/dt* of SSs is 530 mV/ms (440 to 540 mV/ms in experiments). The *d^2^v/dt^2^* of SSs shows a biphasic increase, with the first component reflecting the contribution of axial current from the AIS and the second one reflecting the contribution of somatic Na^+^ current. The peak value of *d^2^v/dt^2^* is 6600 mV/ms^2^ (5600 to 8200 mV/ms^2^ in experiments).

Due to the absence of dendritic Na^+^ channels (Stuart and Hausser, 1994) and the large impedance load of elaborate dendritic trees (Vetter et al., 2001), SSs fail to invade the dendritic tree (Llinas and Sugimori, 1980). The distance-dependent decay of SS amplitudes in the model matches experimental data (Fig. 1E; also close to Stuart and Hausser (1994)). The model also performs well in response to somatic current injections. The F-I curve falls within the experimentally measured range (Fig. 1F). Similar with Fernandez et al. (2007), our model has type-I excitability. When a large enough negative current is injected, the cell is silenced and shows a typical ‘sag’ response (illustrated in the inset of Fig. 1G). Both the peak and steady state of the ‘sag’ response are within the range of experimental data (Fig. 1G).

**Figure 1.**
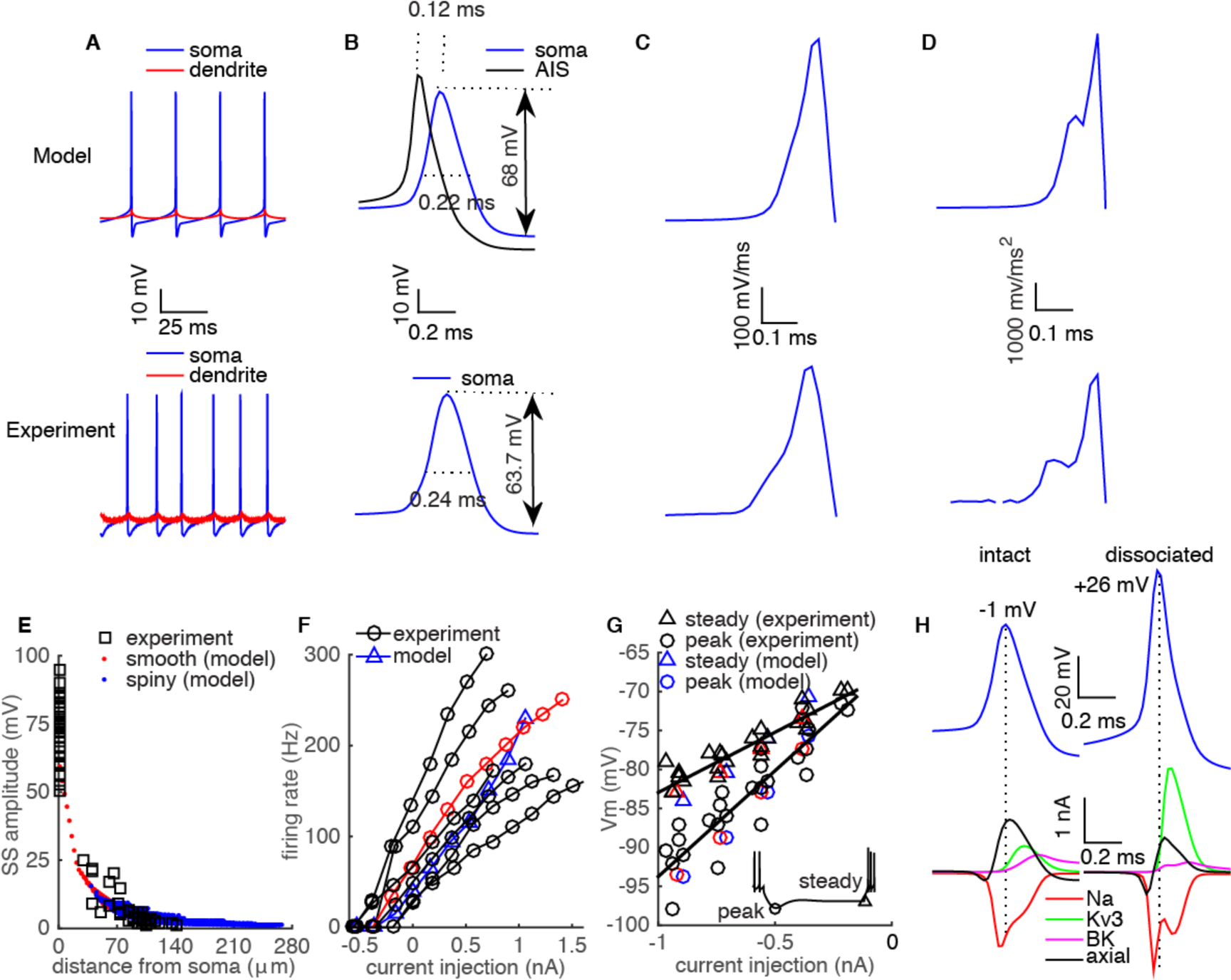
Model properties compare well with experimental data. **A**. Spontaneously firing SSs and dendritic membrane potentials (103 μm from soma) from the model and experiments (dendritic site is ~ 100 μm from soma). **B.** A single SS from the model and experiments. The spike at the AIS is aligned to measure the axosomatic delay in the model. The *dv/dt* and *d^2^v/dt^2^* of the SS in the model are compared with experimental data in **C** and **D** separately. **E**, Distance-dependent decay of SS amplitudes in the model and experimental data (Ohtsuki et al. (2012)). **F**, The F-I curve of the model compared with experimental data. **G**, The peak and steady state values of ‘sag’ responses caused by negative current injections are compared between model and experimental data. In F-G, red symbols are from the example cell used in A-D. The data used in A-D, F-G are from Rancz and Hausser (2010). **H**, Distinct spike peaks and principal ionic current profiles in the intact and the dissociated PC model. In the dissociated PC, only the dendritic root section before the first bifurcation is preserved. Ionic currents are computed at the central segment (surface area of 204 μm^2^) of the soma.

This good overall performance in response to current injections suggests that the model is reliable in predicting excitatory and inhibitory synaptic responses in PCs.

Dissociated PCs are often used to investigate the role of ionic currents in PC electrophysiological properties (Carter and Bean, 2009; Khaliq et al., 2003; Swensen and Bean, 2003). However, we find the profiles of Na^+^ current (principal depolarization current (Carter and Bean, 2009)) and of Kv3 current (principal repolarization current (Martina et al., 2007)) in an intact PC strongly differ from in a dissociated PC owing to their distinct spike properties (Fig. 1H). Similar with Bekkers and Hausser (2007), the intact PC model exhibits a higher spike threshold, smaller afterhyperpolarization and lower spike peak compared with the dissociated PC. The lower spike peak (−1 mV in the model) compared with the dissociated PC (+26 mV in the model, +32 mV in Carter and Bean (2011) and ~ +40 mV in Bekkers and Hausser (2007)) leads to a much smaller fraction of Kv3 channels activated, due to their high activation threshold (Martina et al., 2007). In the intact PC, the dendrite repolarizes somatic SSs together with the ‘less’ activated Kv3 current. This large axial current (flow into dendrite) keeps pace with SS depolarization to constrain its peak to -1 mV (a smaller axial current exists in the dissociated PC due to the remaining dendritic stump). At the peak of the spike, the ‘dip’ in the Na^+^ current observed in dissociated PCs (Carter and Bean, 2009) is absent in the intact PC model.

The Na^+^ entry ratio (the ratio of total Na^+^ influx to the Na^+^ influx during the rising phase (Carter and Bean, 2009)) can decrease from 3.0 in the dissociated PC model to 2.2 in the intact PC model. Although dendrites lack Na^+^ channels, SSs passively propagate into dendrites and partially activate Ca^2+^ channels in proximal dendrites. Therefore, we compute the ATP molecules required for each SS to better understand their energy consumption (Attwell and Laughlin, 2001). In our model, 4.3 x10^7^ ATP molecules are required to pump Na^+^ ions and Ca^2+^ ions out of the PC after a SS, with 60% of the energy expended on Na^+^ ions.

### CF responses in the PC

When PCs receive synaptic inputs from CFs, stereotypical CSs occur at their somas. CF responses at different sites of a spontaneously firing PC model are shown in Fig. 2A. The peak *dv/dt* of the first spikelet in the CS is ~ 270 mV/ms larger than that of the SS, as confirmed by *in vitro* experimental data (Fig. 2B). The large peak *dv/dt* of the first spikelet ensures its reliable propagation down to the cerebellar nuclei (CN) (Khaliq and Raman, 2005; Monsivais et al., 2005). The CS still initiates first at the AIS (Palmer et al., 2010) with an axosomatic delay of the first spikelet of 0.06 ms, which is much shorter than that of the SS (Fig. 2C). This decreased axosomatic delay is caused by the depolarizing CF EPSC during the initial depolarization phase (Fig. 2D). The model also replicates varied CF responses due to bursting CF input (Mathy et al., 2009) (Fig. S1).

In agreement with previous experimental findings (Swensen and Bean, 2003), the SK2 current terminates the CS (Fig. 2D). Each spikelet of the somatic CS in our model, except the last one, is sharp (Fig. 2A), suggesting they are generated by Na^+^ channels (Fig. 2D). Two factors determine the absence of Ca^2+^ spikes at the soma. First, the P-type Ca^2+^ current is much smaller than the Na^+^ current; therefore, the Na^+^ current shunts it. Second, K^+^ currents are always larger than the P-type Ca^2+^ current (Fig. 2D). Magnitudes of the CF EPSC and the net axial current are comparable to somatic ionic currents (Fig. 2D), suggesting an important role in generating somatic CSs. Consequently, a variation of either CF EPSC or axial current has the potential to alter somatic CSs, as shown in later sections.

The dendritic root section (the section between the soma and the first dendritic bifurcation) is electrically compact with the soma and shows mainly a passive propagation of a somatic CS. With distance from the soma, the clamping effect of the somatic CS on the dendrite weakens. The P-type Ca^2+^ current gradually dominates and cause Ca^2+^-driven dendritic responses in the PC dendrite (Fig. 2A, E-F).

**Figure 2.**
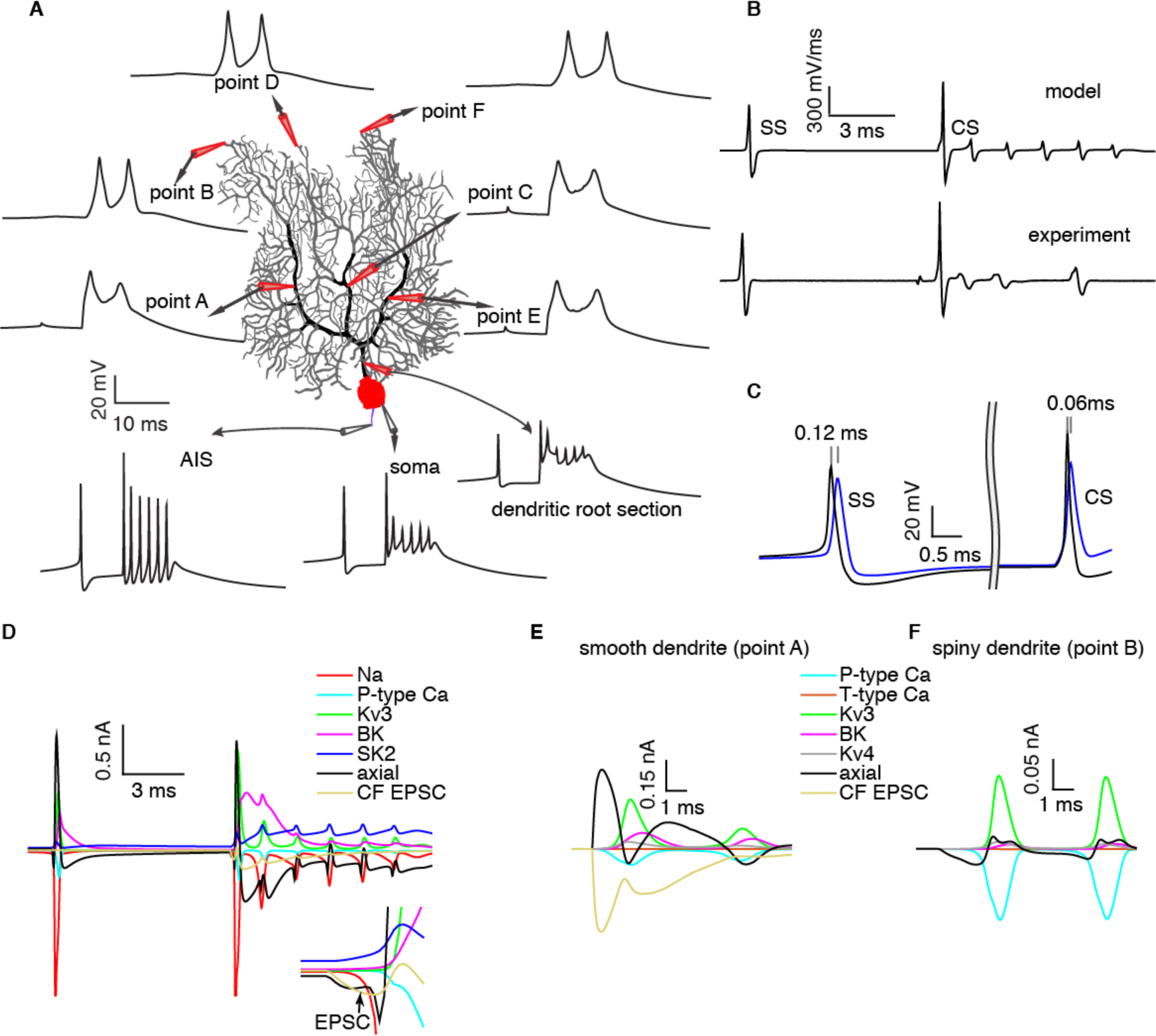
The generation of CF responses in the PC. **A.** CF responses at different sites in the PC model. Two sites on each main branch are selected. Points A (smooth dendrite) and B (spiny dendrite) on branch 1; Points C and D on branch 2; Points E and F on branch 3. CF responses at dendritic root section, soma and AIS are labeled by corresponding arrows. **B.** The peak *dv/dt* of the first CS spikelet increases compared with a SS in both model and experiment (Warnaar et al. (2015). **C.** The axosomatic delay of the first spikelet in the CS decreases compared with a SS. **D.** The composition of the somatic CS. Currents at the initial depolarization phase are enlarged in the inset. Dendritic currents at points A and B are shown in **E** and **F** in sequence. The surface areas of the segments chosen at the soma, point A and point B are 204 μm^2^, 64 μm^2^, and 23 μm^2^, respectively. CF EPSC is absent in the spiny dendrite. Notice different scales used in D-F due to different segment surface areas.

### Factors regulating the initiation and propagation of dendritic spikes

Here we systematically explore how different voltage states, caused by varying the somatic holding current, regulate CF-evoked dendritic responses and analyze the biophysical mechanisms. We find depolarization facilitates higher peaks of dendritic responses with further propagation into distal spiny dendrites. In distal spiny dendrites, axial currents from proximal smooth dendrites are the only current sources that can depolarize to approach the Ca^2+^ spike threshold and trigger a dendritic spike (Fig. 3J-L). When the soma is held at -76 mV (Fig. 3A), the A-type Kv4 current is larger than the P-type Ca^2+^ current in spiny dendrites (Fig. 3J). Consequently, axial currents cause only small passive depolarizations in spiny dendrites (Fig. 3D), and P-type Ca^2+^ channels are hardly activated (Fig. 3J). In smooth dendrites, the CF provides powerful synaptic input to depolarize (Fig. 3D). However, this depolarization is still passive because the Kv4 current and axial currents (flow into distal parts) stop the P-type Ca^2+^ current from dominating depolarization (Fig. 3M). Accordingly, the dendritic voltage responses decrease with distance from the soma in the whole dendrite (Fig. 3G). Distance-dependent propagation of voltage responses matches experimental recordings under the same conditions (Fig. 3G). As the soma is held at -70 mV (Fig. 3B), dendritic spikes occur in part of the dendritic tree (the left half, proximal to the soma), where the voltage responses now increase with distance from the soma. However, the responses still decay with distance from the soma in other parts of the dendritic tree (Fig. 3B,E,H). With depolarization of the soma to -61 mV (Fig. 3C), the Kv4 current gradually inactivates and becomes smaller than the P-type Ca^2+^ current in both smooth and spiny dendrites during the initial depolarization phase (Fig. 3L,O). As a result, the P-type Ca^2+^ current depolarizes the dendrites and activates more Ca^2+^ channels in a positive feedback loop until the Kv3 current is highly activated to repolarize the spikes. Dendritic spikes occur globally and their peaks increase with distance from the soma (Fig. 3F,I). In line with voltage responses, the dendritic Ca^2+^ influx is also facilitated by depolarization, causing larger and more widespread elevations in dendritic Ca^2+^ concentration (Fig. 3P-R). Additional CF responses at other voltage states are shown in Fig. S2.

**Figure 3.**
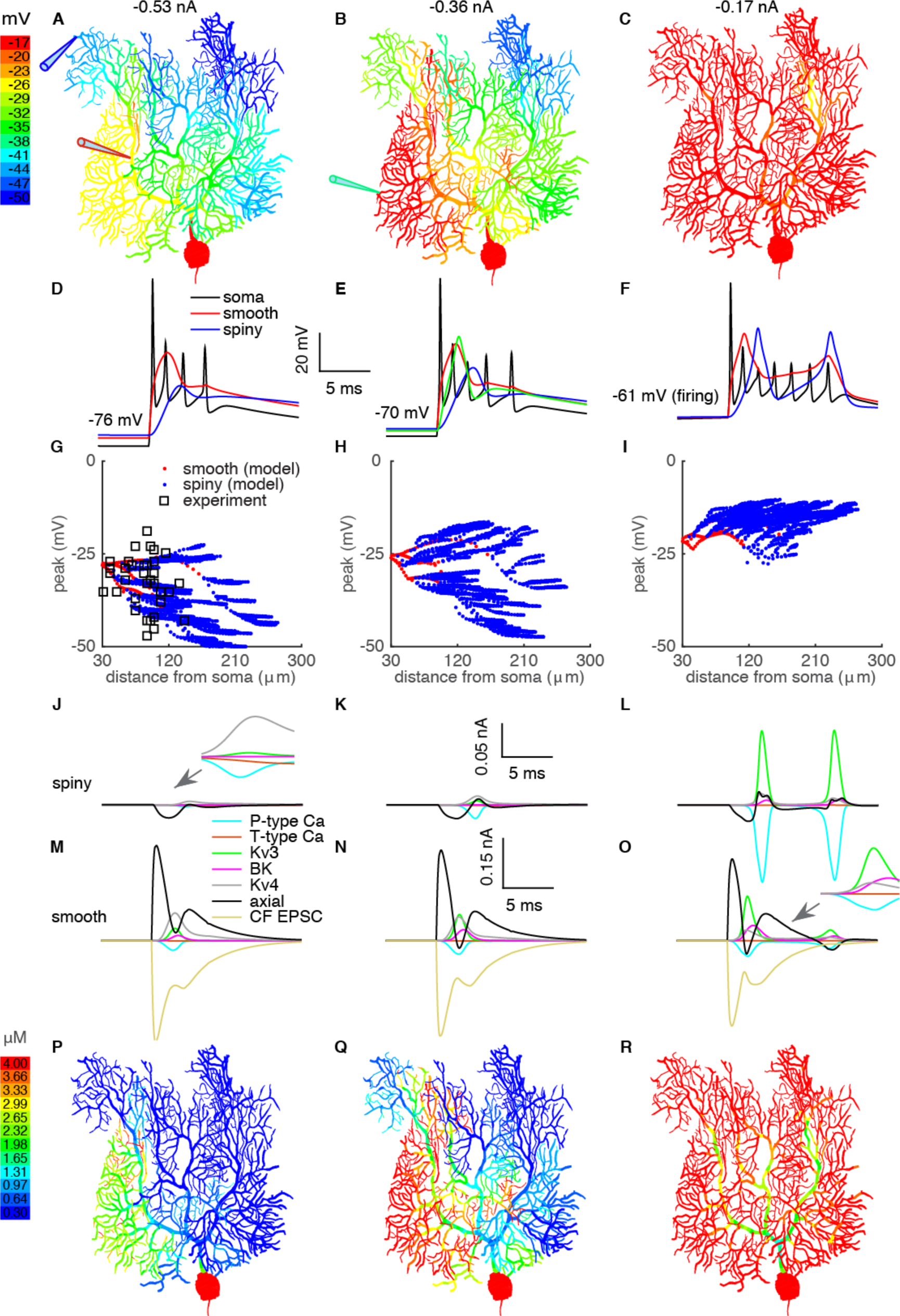
Voltage states regulate CF-evoked dendritic spike generation and propagation. From left to right column, the holding potentials are -76 mV, -70 mV and -60 mV, respectively. **A-C**. Color-coded voltage responses. **D-F**. Somatic CSs, voltage responses on the smooth dendrite and spiny dendrite. In **E**, an extra example voltage response (green trace, position indicated in B) on the spiny dendrite is shown. **G-I**. Distance-dependent propagation of voltage responses. Experimental data in G are from Ohtsuki et al. (2012). **J-L** and **M-O** show ionic currents in the spiny dendrite and smooth dendrite respectively. The surface areas of the segments chosen at the smooth dendrite and spiny dendrite are 64 μm^2^ and 23 μm^2^ respectively. **P-R**. Color-coded Ca^2+^ concentrations.

The critical role of Kv4 current in regulating dendritic spike propagation is further demonstrated in Fig. S3. After blocking dendritic Kv4 current, dendritic spikes reliably propagate to the whole dendrite at more hyperpolarized conditions. We further test whether other K^+^ currents regulate the propagation of CF-evoked dendritic response. Blocking the high threshold-activated Kv3 current surprisingly facilitates the propagation of dendritic spikes (Fig. S4) (Otsu et al., 2014; Zagha et al., 2010). Unlike the ‘brake’ mechanism of Kv4 current, dendritic Kv3 current mainly narrows the CF response and only activates when the response is large enough (Fig. 3 J-O). Blocking the Kv3 current broadens the dendritic responses at smooth dendrites and facilitates their propagation to spiny dendrites. The effect of broadening dendritic responses on their propagation is further demonstrated in Fig. S5. Dendritic BK current does not significantly affect the propagation of CF-evoked dendritic responses (data not shown).

Taken together, these results suggest that availability of dendritic K^+^ channels not only regulates the amplitude (Kitamura and Hausser, 2011; Rokni et al., 2009) but also gates the spatial range of Ca^2+^ influx (Ohtsuki et al., 2012; Otsu et al., 2014; Zagha et al., 2010). Voltage conditions can also explain the distinct dendritic spike waveforms recorded *in vitro* by different groups (Davie et al., 2008; Ohtsuki et al., 2012) in Fig. S6.

### Inhomogeneous excitability of the dendritic tree

Similar to the first dendritic spikelet (Fig. 3B), the secondary dendritic spikelet occurs asynchronously in different branches around its threshold (Fig. 4). It initially occurs at branch 1; then it gradually occurs at branch 2 and finally occurs at branch 3 with slightly increasing depolarization. Even when secondary dendritic spikelets occur globally, their time to peak still differs between branches (Fig. 4C). They peak at 8.36 ms, 10.74 ms and 13.64 ms at points A, C and E respectively (relative to CF activation time). With further depolarization, secondary dendritic spikelets peak more synchronously (with 0.89 nA current injection, the respective values become 6.06 ms, 6.88 ms and 7.32 ms). The asynchronous occurrence of individual spikelets reflects the inhomogeneous excitability of each dendritic branch. In the model, the density of ionic currents is homogeneous in the spiny dendritic tree. Therefore, morphological differences between individual branches may account for heterogeneous excitability. The excitability of each branch is determined by the ratio of spiny dendrite area to smooth dendrite length in that branch (essentially the ratio of spiny dendrite capacitance load to CF synaptic input, ‘load/input’ ratio, see Methods). The ‘load/input’ ratios are 69, 96 and 130 at branches 1, 2 and 3 respectively, explaining their heterogeneous excitability. Moreover, within each ‘big’ branch, ‘small’ child branchlets also show heterogeneity in excitability (Fig. 3A-C), which are caused by other morphological factors such as the electrotonic distance to smooth dendrites.

**Figure 4.**
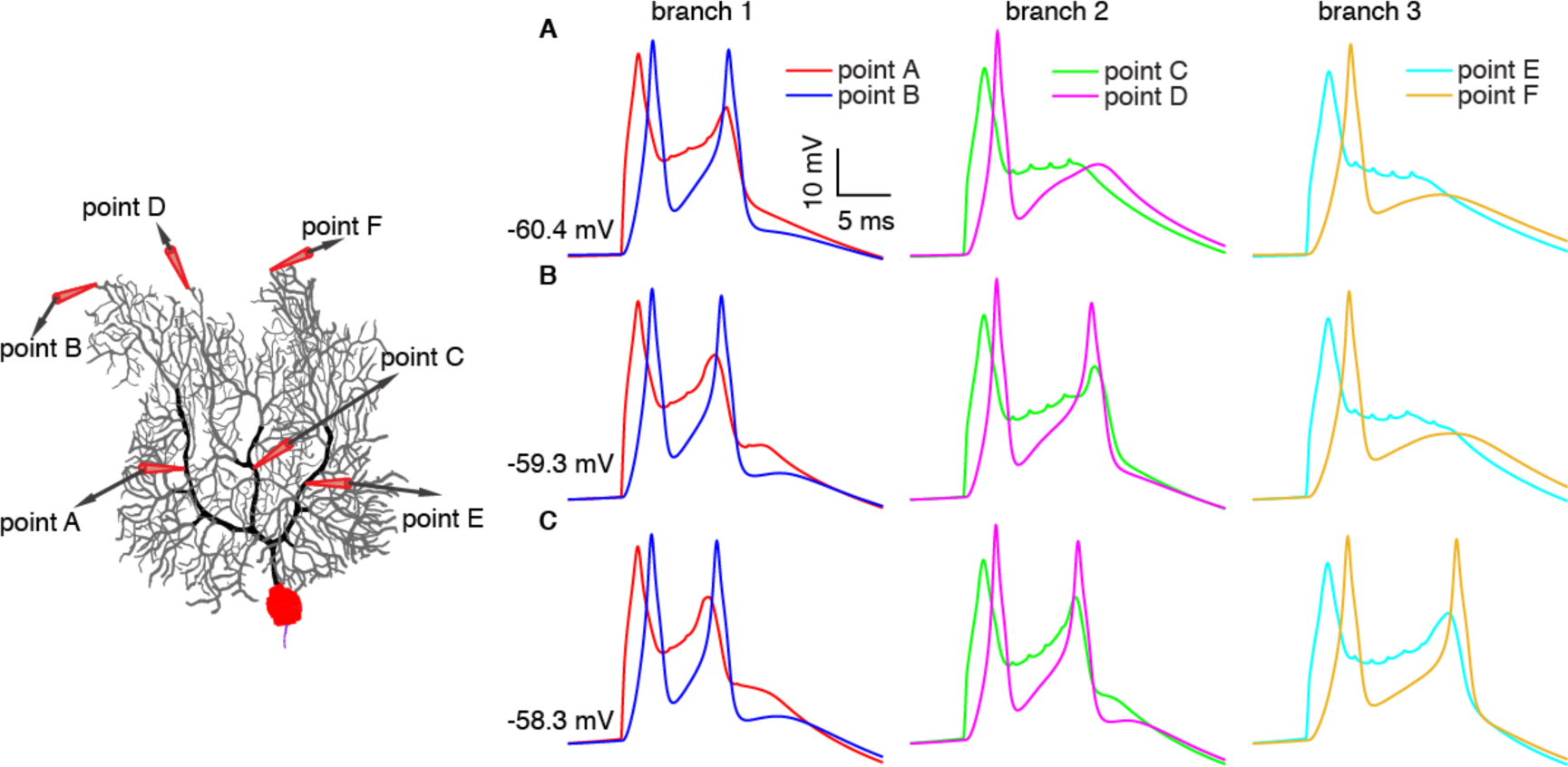
Branch-specific occurrence of secondary dendritic spikelets. In all simulations, the holding current is -0.17 nA. CF input is activated at three slightly different dendritic voltage states (different phases of somatic interspike intervals) as shown in **A, B**, and **C**.

K^+^ currents activated near ‘resting’ dendritic membrane potentials may enhance the heterogeneous excitability. The whole dendrite becomes more homogeneous at hyperpolarized levels after blocking the Kv4 current (Fig. S3). Essentially, uneven distributed ‘load/input’ ratios underlie the heterogeneous dendritic excitability, but dendritic Kv4 channels enhance it.

### Voltage- and phase-dependent somatic CSs

Similar with dendritic responses, somatic CSs also vary significantly with depolarization (Fig. 5A-C). The spikelet numbers in CSs increase with initial depolarization, but decrease with further depolarization. A somatic CS results from the interaction of intrinsic ionic currents, axial current, and CF EPSC (Fig. 2D). At low voltage, although Na^+^ channel availability decreases slightly (Fig. 5C), increased axial current from larger dendritic responses (Fig. 5A) dominates and triggers more somatic spikelets. Interestingly, the largest number of somatic spikelets coincides with the longest occurrence latency of secondary dendritic spikelets (Fig. 5A). With further depolarization, Na^+^ channel availability decreases at a higher rate (Fig. 5C). Also, secondary dendritic spikelets peak faster and closer to a preceding somatic spikelet (Fig. 5A). The shortened interval between secondary dendritic spikelets and their preceding somatic spikelet means fewer Na^+^ channels are recovered to evoke an extra somatic spikelet (right axis of Fig. 5D). The critical role of secondary dendritic spikelet timing in evoking an extra somatic spikelet agrees with previous experimental observations (Davie et al., 2008). Both factors decrease somatic spikelets with further depolarization. Concurrent with spikelet numbers, CS durations are regulated by voltage states, as represented by the timing of the last spikelets (Fig. 5B).

**Figure 5.**
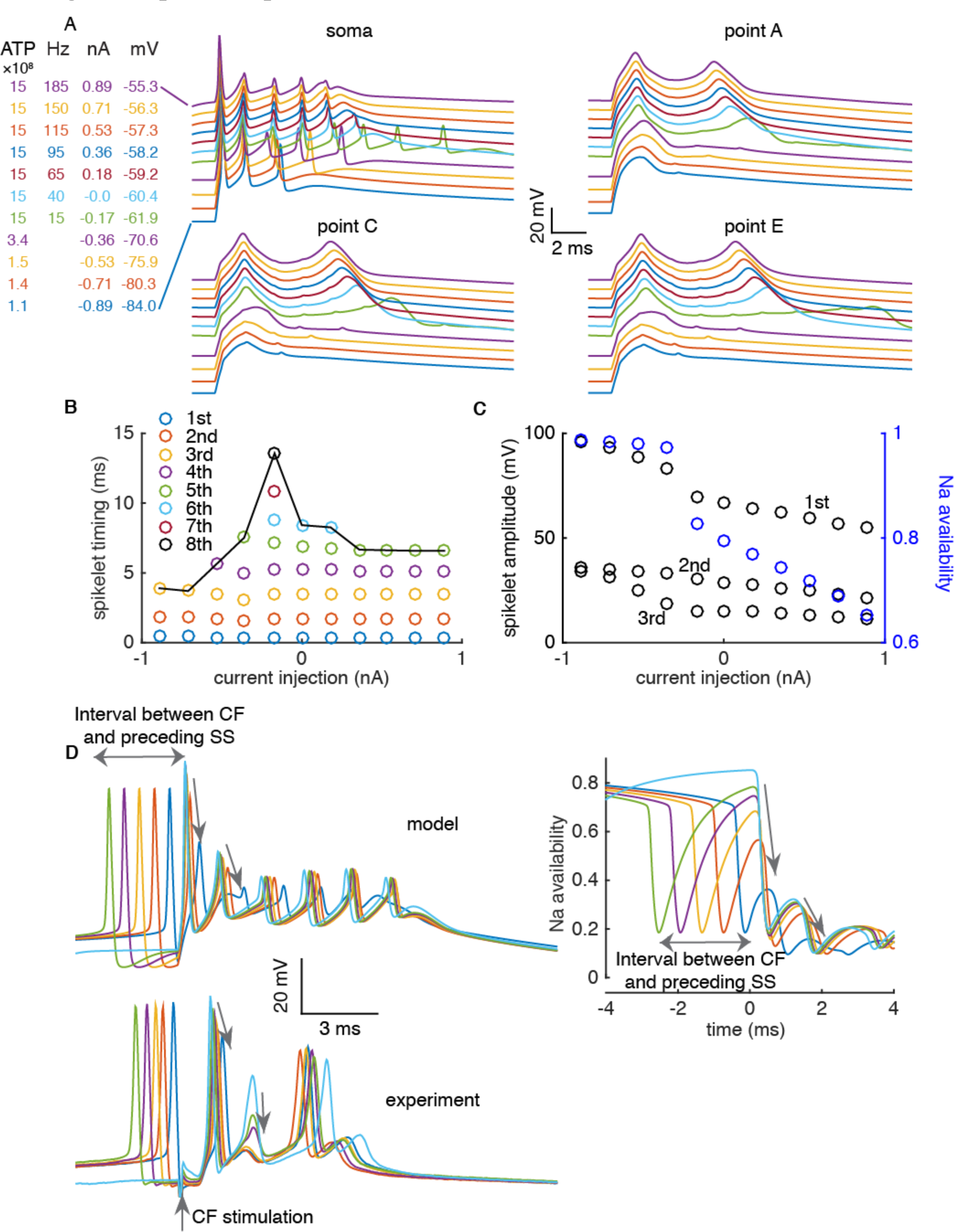
Voltage- and phase-dependent somatic CSs. **A**. Voltage states affect somatic CSs and corresponding dendritic responses. The definition of dendritic sites is the same as in Fig. 2. The holding currents, the FRs and the consumed ATP molecules at each voltage state are listed on the left. **B**. Voltage states modulate CS durations, represented by the timing of the last spikelets. **C**. In parallel with Na^+^ channels availability (blue circles), amplitudes of the first three spikelets in the CS (black circles) decrease with depolarization. **D**. Amplitudes of the first and second spikelets are phase-dependent in model and experiments (Warnaar et al. (2015)). Phase-dependent recovery of Na^+^ channels is shown in upper right panel.

With depolarization, the consumed ATP molecules during CSs increase from 1.1 × 10^8^ (−84 mV) to 1.5 x10^9^ for a CS with two dendritic spikelets (nearly 40 times as large as in a SS), with 96% of the energy expended on Ca^2+^ ions (Fig. 5A). The increased ATP consumption with depolarization is due to increased Ca^2+^ influx (Fig. 3,5A,S2), but relatively insensitive to somatic spikelet variations.

Voltage states also affect spikelet amplitudes in CSs. We analyzed the first three spikelets, which exist in all simulated voltage conditions. Spikelet amplitudes decrease with depolarization due to reduced availability of Na^+^ channels (Fig. 5C). Individual spikelet amplitude is critical because it determines the probability of propagation down to the CN (Khaliq and Raman, 2005; Monsivais et al., 2005).

Finally, we observed phase-dependency of somatic CSs. Amplitudes of the first two spikelets in CSs decrease and the spikelets become more blunted with shortening of the interval between the CF activation and its preceding SS (Fig. 5D). This phase-dependency reflects the time course of Na^+^ channel recovery from inactivation. We confirmed this phase-dependency by reanalyzing *in vitro* experimental recordings.

So far, voltage states of the PC model were manipulated by varying somatic holding currents. We implemented more realistic *in vivo* simulations (Jelitai et al., 2016), in which the net balance of excitatory PF and inhibitory synaptic inputs determines the dendritic voltage states (Fig. S7). We find that the effect of voltage states on CF responses found *in vitro* also holds *in vivo.* We also analyze pauses following CSs (data *in vitro* not shown) and find their durations decrease with depolarization in our model (Fig. S7I), due to the larger depolarization force.

### Spatially constrained modulation by clustered PF/stellate cell synaptic input

Sensory stimuli can activate clustered PF synaptic inputs (Wilms and Hausser, 2015). Here we examine how simultaneous clustered PF or stellate cell synaptic inputs regulate CF responses. We find that clustered synaptic inputs regulate dendritic responses in a spatially constrained manner. A secondary dendritic spikelet only occurs at branch 1 with CF input in isolation (Fig. 6A). With 5 PF synapses simultaneously activated at the indicated branchlet of branch 2, the secondary dendritic spikelet also occurs at this branchlet. Concurrently, an extra spikelet is evoked in the somatic CS. Nonetheless, clustered PF synaptic inputs within branch 2 don’t significantly affect other branches. Seemingly paradoxically, the extra somatic spikelet is eliminated if the number of simultaneously activated PF synapses increases to 20. As explained previously, Na^+^ channels need time to recover from inactivation after each somatic spikelet. Increasing PF synaptic inputs accelerates secondary dendritic spikelets, making them occur closer to the preceding somatic spikelet, and reducing the likelihood of triggering an extra spikelet due to fewer recovered Na^+^ channels. Similarly, simultaneous activation of 10 stellate cell synaptic inputs can evoke an extra somatic spikelet by delaying the secondary dendritic spikelets at branch 2 (Fig. 6B).

**Figure 6.**
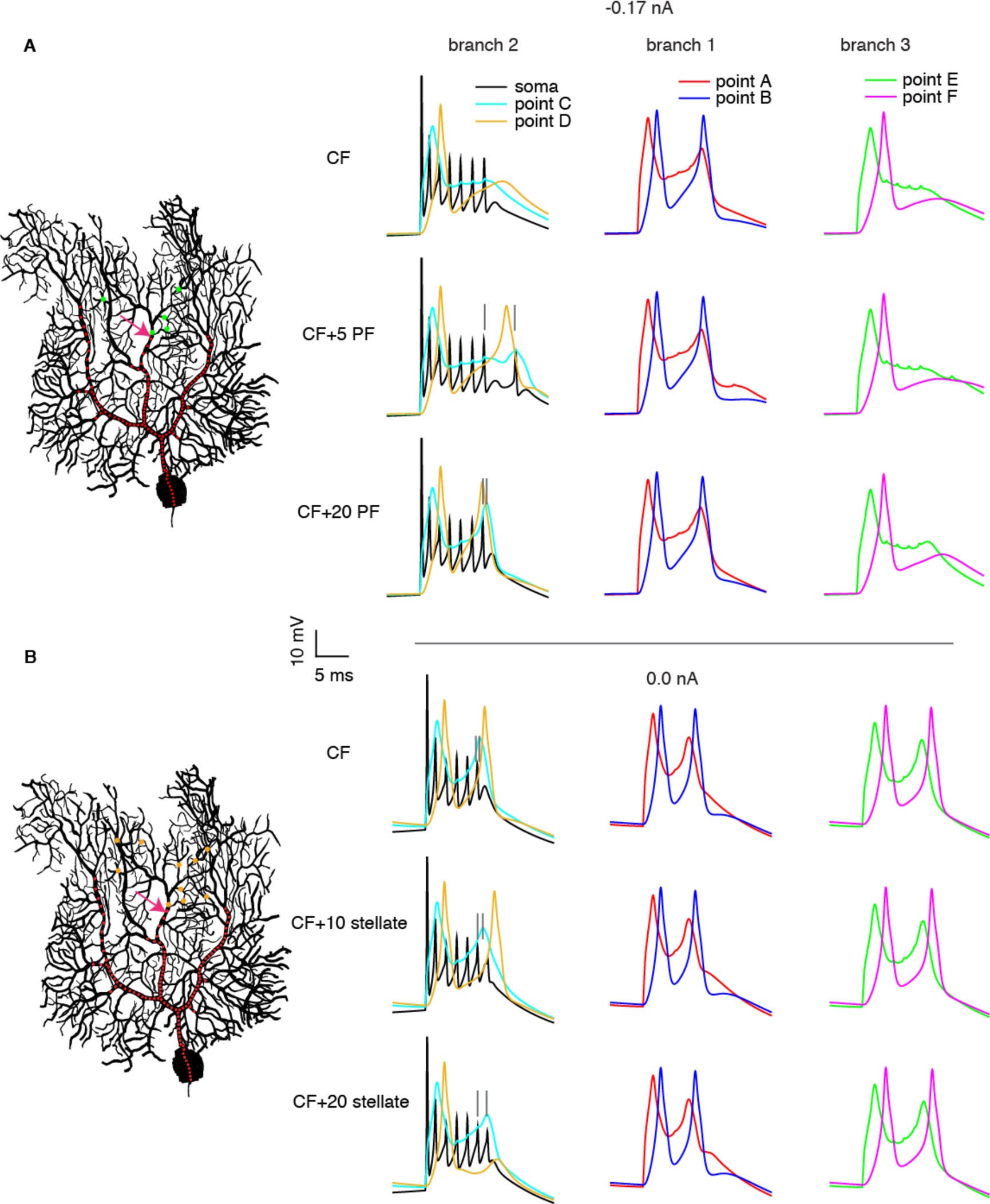
Simultaneous clustered PF or stellate cell synaptic input regulates the CF-evoked somatic CSs by locally modulating the dendritic responses. **A**. Somatic and dendritic responses with only CF input (red dots), CF input + 5 PF synaptic inputs and CF input + 20 PF synaptic inputs are shown from top to bottom. PF synapses are randomly distributed on the child branchlet of the indicated dendritic section (green dots). **B**. Somatic and dendritic responses with only CF input, CF input + 10 stellate cell synaptic inputs and CF input + 20 stellate cell synaptic inputs are shown from top to bottom. Similar placement of stellate cell synapses (orange dots). The definition of recorded dendritic sites is the same as in Fig. 2.

Again, dendritic responses at other branches are minimally affected. The secondary dendritic spikelet still occurs in the proximal smooth dendrite even when it is eliminated in the distal part by activating 20 stellate cell synaptic inputs.

We also show that compartment-specific dendritic excitability (Ohtsuki et al., 2012), simulated here by regional Kv4 block, modulates dendritic responses in a spatially precise way (Fig. S8).

### Paired-pulse depression regulates CF responses in the PC

CFs can transiently fire at ~ 5 - 8 Hz after sensory stimuli (Najafi et al., 2014; Warnaar et al., 2015). Paired-pulse depression (PPD) has been demonstrated for consecutive CF synaptic inputs *in vitro* (Andjus et al., 2005; Hansel and Linden, 2000) and it may support the correlation between instantaneous CF FRs and varied CS spikelet numbers. To investigate the effect of PPD on CF responses, we systematically explore CF responses with baseline or 20% depressed CF EPSC (~ 4 Hz of instantaneous CF FR in experiments by Andjus et al. (2005)). We find depressed CF inputs consistently reduce dendritic responses. At low voltage, the peak of the dendritic response decreases (Fig. 7A); at higher voltage, the appearance of a secondary dendritic spikelet is prevented (Fig. 7B) or delayed (Fig. 7C). The spikelet numbers in somatic CSs show no changes, decreases or increases with depressed CF inputs at different voltages (Fig. 7A-C). A depressed CF input always delays individual somatic spikelets by providing less depolarization current. At low voltage, 20% depression of the CF EPSC and slightly reduced dendritic response are insufficient to vary somatic spikelet number and only increase CS duration (Fig. 7A and Case I in Fig. 7D). At medium voltage, baseline CF input evokes a secondary dendritic spikelet. The increased axial current (the later stage of ionic currents in Fig. 7B) triggers extra somatic spikelets due to the relatively large availability of Na^+^ channels (see Na^+^ current amplitude). Therefore, by not causing a secondary dendritic spikelet, depressed CF input decreases CS duration because of fewer somatic spikelets, although they are delayed (Fig. 7B and Case II in Fig. 7D). At high voltage, depressed CF input delays individual spikelets and allows for larger recovery of Na^+^ channels. The delayed secondary dendritic spikelet also facilitates occurrence of extra somatic spikelets. Thus, depressed CF input increases CS duration (Fig. 7C and case III in Fig. 7D).

**Figure 7.**
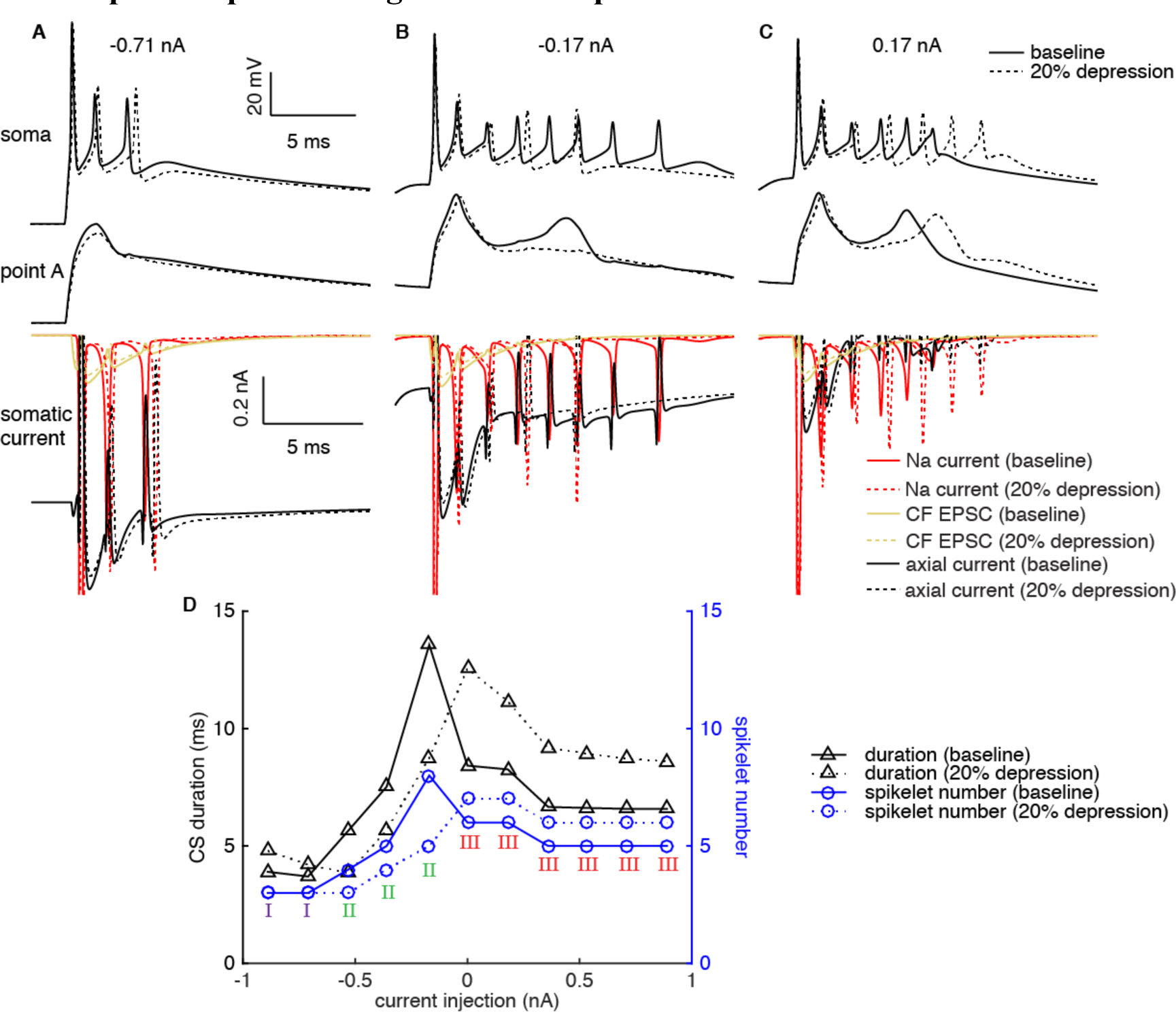
PPD regulates CF responses. CF responses (somatic and dendritic) and somatic ionic currents with unchanged (case I), decreased (case II) and increased (case III) spikelet numbers per CS by depressed CF EPSC are shown in **A, B** and **C** in sequence. Ionic currents are computed in the central segment of the soma and Na^+^ currents during the first spikelet are truncated. **D**. The effect of depressed CF EPSC on the spikelet number and duration of the CSs.

Voltage ranges for the occurrence of cases I-III depend on the degree of depression. For example, when CF EPSC is depressed by 10%, case I can occur at both low and high voltages (data not shown). Also, when CF FRs are higher, incomplete recovery of ionic currents from preceding CS such as SK2 and the dendritic hyperpolarization may vary the posterior CS in a more complicated way.

PPD can bridge the gap between instantaneous CF FRs and CS variations. Our results reconcile the observed unchanged (Warnaar et al., 2015), decreased (Khaliq and Raman, 2005; Maruta et al., 2007), and increased (Burroughs et al., 2017) somatic spikelet numbers at higher CF FRs.

## Discussion

We built a new PC model based on extensive experimental data to systematically investigate CF responses. Simulation results help to reveal the mechanisms causing spikes on different parts of PCs, reconcile conflicting experimental data and make new predictions.

### Model accuracy

We constrained model parameters by hand tuning and tried to replicate as many experimental observations as possible. We started by constraining somatic ionic currents based on ionic current measurements taken from dissociated PCs. These data allowed us to uniquely constrain principal somatic currents. Subsequently, the ionic currents in the AIS and dendrites were constrained to obtain the spike properties summarized in Fig. 1. It was more difficult to constrain dendritic ionic currents due to insufficient data. The total Ca^2+^ influx during one dendritic spikelet was constrained by the experimentally estimated value. In addition, the replicated CF-evoked dendritic responses under different conditions (Fig. 3,S2,S4,S6) suggest that the dendrite model produces realistic results. Nonetheless, there is still room for improvement in the future. Due to the lack of specific data, we did not try to reproduce the role of dendritic SK2 current (Ohtsuki et al., 2012; Womack and Khodakhah, 2003) and Kv1 current (Khavandgar et al., 2005) in regulating dendritic excitability. These currents may further enhance the heterogenous excitability and regulate dendritic responses. The data we used to directly validate our simulation results are all from *in vitro*. We hope our study will inspire experiments to validate modifiable CF responses *in vivo.*

Our PC model follows the tradition of the biophysical modeling by De Schutter and Bower (1994a, 1994b), which has successfully predicted experimental findings (Steuber et al., 2007). Although the 1994 model is outdated by recent knowledge of ionic channels in PCs, there was no suitable replacement until now. A PC model by Khaliq et al. (2003) has no dendrite and therefore can’t be used to simulate synaptic responses. A recent PC model (Masoli et al., 2015) lacks extensive validation against experimental data. For example, it fails to produce the stereotypical ‘sag’ responses to hyperpolarization.

### Physiological predictions and implications

The model allows us to predict the metabolic cost of spiking in PCs. In a cortical pyramidal neuron, ~ 4* 10^8^ ATP molecules are consumed to restore the Na^+^ and K^+^ ion gradients after each spike (Attwell and Laughlin, 2001). Carter and Bean (2009) reported that spiking is energetically less efficient in PCs compared with cortical pyramidal neurons. However, their study was performed in dissociated PCs. We find the intact PC has a smaller Na^+^ entry ratio compared with the dissociated PC (Fig. 1H), suggesting that dendrites increase the metabolic efficiency of somatic spikes. Given the energetic cost of spikes, it will be interesting to explore whether this dendritic role also applies to other neurons. Compared with Na^+^ entry ratio, the estimated energy consumption may be a better measure because of the Ca^2+^ channels distributed in soma, axon and elaborate dendritic tree. Interestingly, although cortical pyramidal neurons have a low Na^+^ entry ratio (~ 1.1 by Carter and Bean (2009)), their estimated energetic cost of a single spike is ~ 8 times larger than a SS in PCs, due to the lack of Na^+^ channels in PC dendrites. Each CS consumes more ATP molecules, which consumes nearly the same amount of energy as 40 SSs during 1 sec in our model. Possibly, previously reported negative correlation (Cerminara and Rawson, 2004) between the FRs of SSs and CSs may be a protective mechanism given the limited energy supply in the cerebellum.

CF-evoked dendritic spikes and somatic CSs are generated by different channels intrinsically, but they communicate via axial currents through the dendritic root section (Fig. 2). Due to technical challenges, dendritic patch-clamp recording is usually performed on smooth dendrites of large diameter. Though we could reproduce the experiment by Davie et al. (2008) in the model (Fig. S9), we show that a single-site dendritic recording can’t reliably reflect spike occurrence and propagation in the whole dendrite. Both the peak and occurrence latency of secondary dendritic spikelets can be modulated. The peak of secondary dendritic spikelets decreases in the somatopetal direction due to unevenly distributed impedances (Roth and Hausser, 2001; Vetter et al., 2001) and K^+^ channels (Martina et al., 2003). When a secondary dendritic spikelet is ‘weak’, with small amplitude and long latency, it may fail to trigger a regenerative dendritic spike in proximal dendrites (branch 1 in Fig. S9). Therefore the role of dendritic spikes in evoking extra somatic spikelets may have been underestimated.

Using the model, we deciphered how voltage-dependent K^+^ currents regulate the amplitude (Kitamura and Hausser, 2011; Rokni et al., 2009) and spatial range of Ca^2+^ influx (Fig. 3,S2-S4,S7) as observed in separate experiments (Ohtsuki et al., 2012; Otsu et al., 2014; Zagha et al., 2010). Vetter et al. (2001) have explored the local propagation of dendritic spikes by exerting an AP clamp at a specific dendritic site. We find that dendritic spikes on the smooth dendrite can overcome the impedance mismatch and propagate to distal dendrites in the absence of the ‘brake’ role of Kv4 in distal dendrites (Fig. S3,S5). Therefore, K^+^ currents activated around the ‘resting’ dendritic membrane potential such as Kv4 in spiny dendrites are necessary to gate the spatial range of CF-evoked dendritic responses. Unlike Kv4 current, Kv3 current narrows the dendritic response to regulate its propagation (Fig. S4,S5). The voltage-related amplitude and spatial range of dendritic responses indicate that PCs can play active roles in response to CF error signals rather than passively accepting instruction from their CF ‘teachers’. PF synapses can undergo not only LTD (with a higher Ca^2+^-induction threshold), but also long-term potentiation (LTP, with a lower Ca^2+^-induction threshold) (Coesmans et al., 2004; Gallimore et al., 2018; Piochon et al., 2016). The voltage-regulated dendritic spike amplitude and spatial range suggest that CF may trigger LTP, a mixture of regional LTP and LTD, or LTD depending on the voltage states of PCs. Such effects will be further modified by stochastic gating of Ca^2+^ and K^+^ channels (Anwar et al., 2013), which was not simulated in this study.

The PC dendritic tree exhibits inhomogeneous excitability in individual branches, implying a computational unit at the level of individual branches, or even smaller branchlets (Fig. 4). Building upon the modulated voltage states of PCs, branch-specific computation can undoubtedly increase the capacity of information processing in PC dendrites, in contrast to the traditional view of the whole dendrite as the computational unit of CF response. Notably, branchlet-specific Ca^2+^ influx has been demonstrated for PF synaptic integration (Wang et al., 2000). When PCs receive concurrent clustered synaptic input (Wilms and Hausser, 2015) or undergo localized dendritic excitability plasticity (Ohtsuki et al., 2012), CF-evoked dendritic responses can be modulated more precisely in space and magnitude to affect the plasticity induction (Fig. 6,S8).

CS shapes are quite variable. In our model, we did not get the ‘small’ second spikelets as observed in some experiments (Davie et al., 2008; Monsivais et al., 2005), but our CS shapes are close to other recordings (Jelitai et al., 2016; Mathy et al., 2009; Otsu et al., 2014). We speculate that CS shapes are cell-dependent. Recently, CS durations have been linked to the degree of trial-over-trial learning (Yang and Lisberger, 2014), with the mechanism underlying variable CSs unresolved. The authors assumed that the variation of CS durations is due to the bursting of CF inputs, based on Mathy et al. (2009). We correlate the spikelet numbers and durations of CSs with voltage states or SSFRs (Fig. 5A-B). Besides reconciling previous contradictory correlations between spikelet numbers and SSFRs (Burroughs et al., 2017; Gilbert, 1976; Khaliq and Raman, 2005; Mano, 1970; Warnaar et al., 2015), our results provide an alternative interpretation of the work by Yang and Lisberger (2014). Our interpretation highlights the importance of all synaptic inputs to the PCs and resulting SSFRs, not just CF, to control learning. Additionally, the CS spikelet amplitudes can be modulated by voltage states (Fig. 5C, S7H). The timing of CF activation (phase-dependency) and concurrent synaptic input also affect somatic CSs (Fig. 5D, 6). All these factors vary CS shapes and may modulate the PC output and affect inhibition of the CN. Although CF-evoked pause decreases at higher firing rates (Fig. S7I), they are very variable (Mathy et al., 2009) when voltage states, CF activation phase or CF input conductance are varied. Sometimes, there is even no ‘obvious’ pause following a CS.

The CF input itself is not ‘all-or-none’ either. The variable number of spikes per CF burst can affect both somatic and dendritic responses (Mathy et al., 2009) (Fig. S1). Furthermore, CF synaptic inputs can be regulated by PPD (Andjus et al., 2005; Hansel and Linden, 2000) and by norepinephrine (Carey and Regehr, 2009). Both of them can depress the CF EPSC to modulate somatic and dendritic responses (Fig. 7). In particular, PPD bridges the gap between instantaneous CF FRs and varied CS spikelet numbers (Burroughs et al., 2017; Khaliq and Raman, 2005; Maruta et al., 2007; Warnaar et al., 2015) and can play an important role in regulating CF responses.

## Methods

All simulations were performed with NEURON, version 7.4 (Hines and Carnevale, 1997). We use the morphology of a 21-day-old Wistar rat PC (Roth and Hausser, 2001). As shown in Fig. 2A, the model consists of an AIS (purple), a soma (red), smooth dendrites (black), and spiny dendrites (gray). The first 17 μm of the reconstructed axon are preserved as the AIS. When exploring the spike initiation site, AIS indicates the end next to the myelinated axon. The model has following passive parameters: Rm = 120.2 kΩ cm^2^, Ri = 120 Ω cm, Cm = 0.64 μF/cm^2^. To compensate for the absence of spines in the reconstructed morphology, the conductance of passive current and Cm are scaled by a factor of 5.3 in the spiny dendrite and 1.2 in the smooth dendrite. The conductance densities of ionic currents in different parts of the model were found by hand tuning. Detailed information about ionic current equations and the process of parameter tuning can be found in Supplementary Information.

The spiking threshold of SSs was defined as the membrane potential at which *dv/dt* = 20 mV/ms. The synaptic conductance activated by CF input is approximated as a biexponential waveform, with 0.3 ms and 3 ms as the rise and decay time constants (Davie et al., 2008). The peak conductance of baseline CF input is 1.2 nS. 500 synaptic inputs are distributed on the soma and smooth dendrite at a constant density per length to simulate the CF input. The conductance varies in the range of 80% - 100% to simulate the EPSC changes by PPD. Due to the elaborate structure, the dendrite is not isopotential with the soma when holding current is injected at the soma. The somatic membrane potential is used to represent the voltage state of the PC except in Fig. 4, in which dendritic membrane potential is used. When measuring the distance-dependence of the peak dendritic response, we selected the first dendritic spikelet if there were two spikelets. To measure the excitability of individual branches, we define a new measure as the ratio of the spiny dendrite capacitance load to CF synaptic input, ‘load/input’ ratio, which can be transformed into ‘area of spiny dendrite’/‘length of smooth dendrite’ within each branch in the model. To exclude the effect of phase-dependence on CSs in Fig. 5, all the CF signals are activated with 3 ms after the preceding SS under different voltage states. To explore how simultaneous clustered synaptic inputs regulate dendritic response and somatic output, PF or stellate cell synaptic inputs are activated 5 ms after the CF input (Fig. 6). To simply mimic the PC voltage states *in vivo*, 1105 PF synapses and 1105 Inhibitory synapses are uniformly distributed on the spiny dendrites. Different voltage states of the PC are achieved by randomly activating excitatory and inhibitory synapses at different average FRs. To estimate the energy cost of spikes, we computed the Na^+^ (3 Na^+^ ions consume 1 ATP molecule) and Ca^2+^ influx (1 Ca^2+^ ion consumes 1 ATP molecule) during a spike in the whole PC (AIS, soma and dendrite) (Attwell and Laughlin, 2001).

## Data availability

The model code will be available from ModelDB.

## Acknowledgements

We thank Ede Rancz, Michael Hausser, Gen Ohtsuski, Christian Hansel, Joao Couto (the data shared by above-mentioned people were used in the manuscript for validation of the simulation results), Bruce Bean, Indira Raman (the data shared by abovementioned people were used to constrain ionic current properties), Mathew Nolan, Edward Zagha, Bernardo Rudy and Paul Mathews for sharing related experimental data for us to understand PC properties.

## Author Contributions

Y.L.Z., S.D., and E.D.S. conceived this study; Y.L.Z. performed all the simulations; Y.L.Z. and E.D.S. analyzed the simulation data; Y.L.Z. and E.D.S. wrote and revised the manuscript.

## Supplementary Information

**Supplementary Figures**

**Figure S1.**
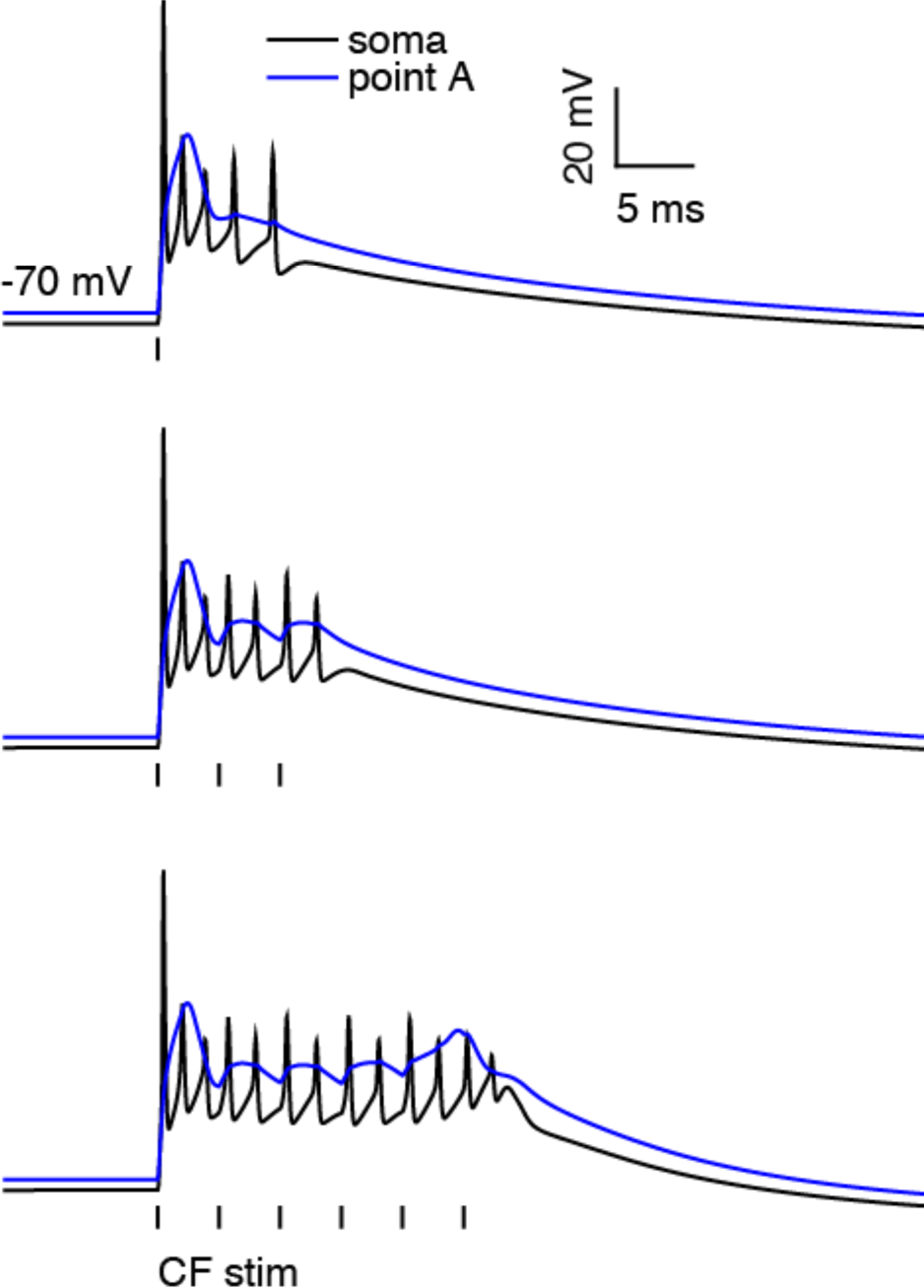
PC responses to CF bursts. Related to Figure 1. We reproduced the experiment by Mathy et al. (2009). Example somatic and dendritic responses to CF burst with 1 (top), 3 (middle) and 6 spikes (bottom) in the CF burst. The simulated instantaneous CF bursting frequency is 250 Hz close to the experimentally reported ~ 273 Hz.

**Figure S2.**
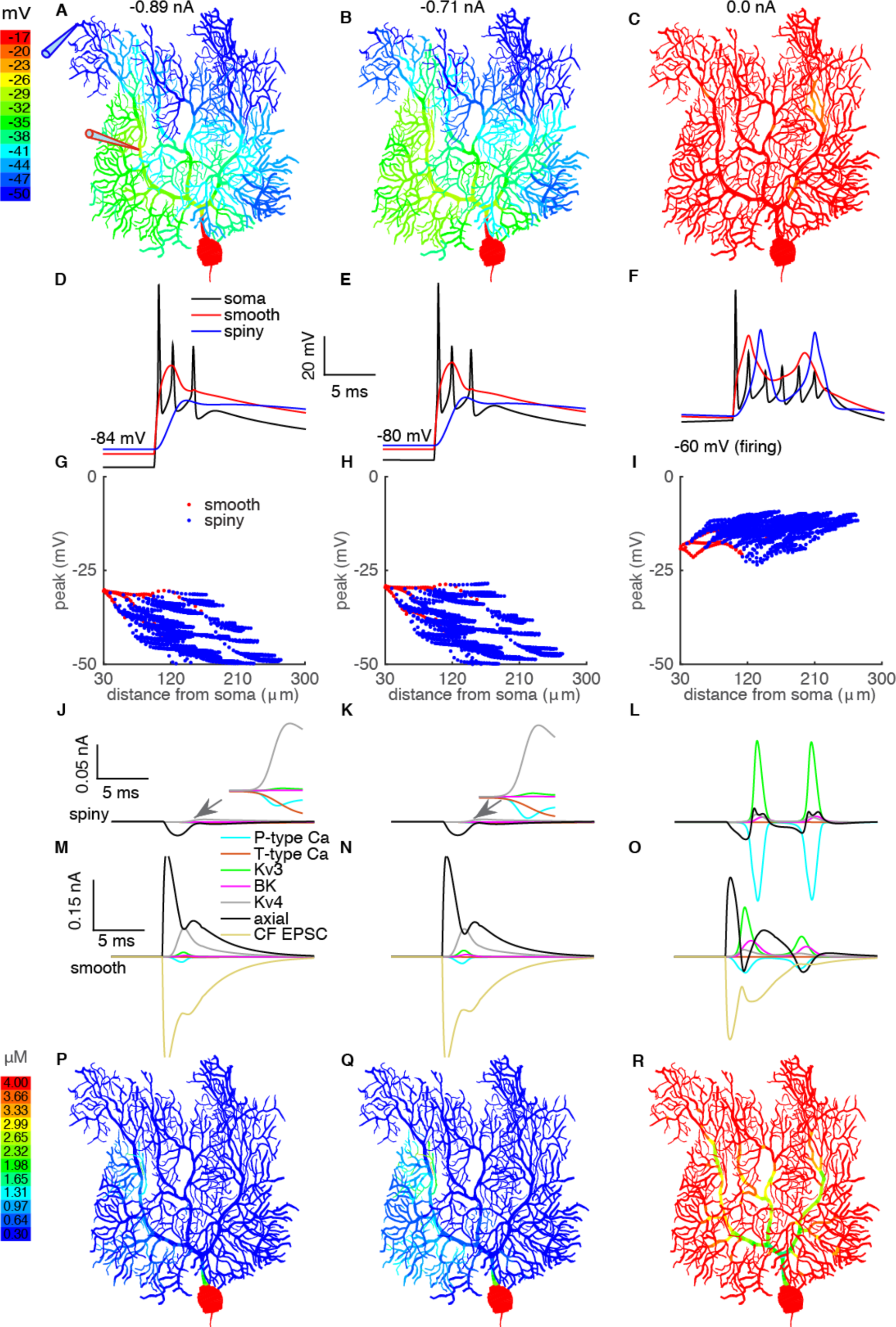
Voltage states regulate CF-evoked dendritic spike generation and propagation. Related to Figure 3. From left to right column, the holding potentials are -84 mV, -80 mV and -60 mV respectively. A-C. Color-coded voltage responses. D-F. Somatic CSs, voltage responses on the smooth dendrite and spiny dendrite. G-I. Distance-dependent propagation of voltage responses. J-L and M-O show ionic currents in the spiny dendrite and smooth dendrite respectively. The surface areas of the segments chosen at the smooth dendrite and spiny dendrite are 64 μm^2^ and 23 μm^2^ respectively. P-R. Color-coded Ca^2^ concentrations.

**Figure S3.**
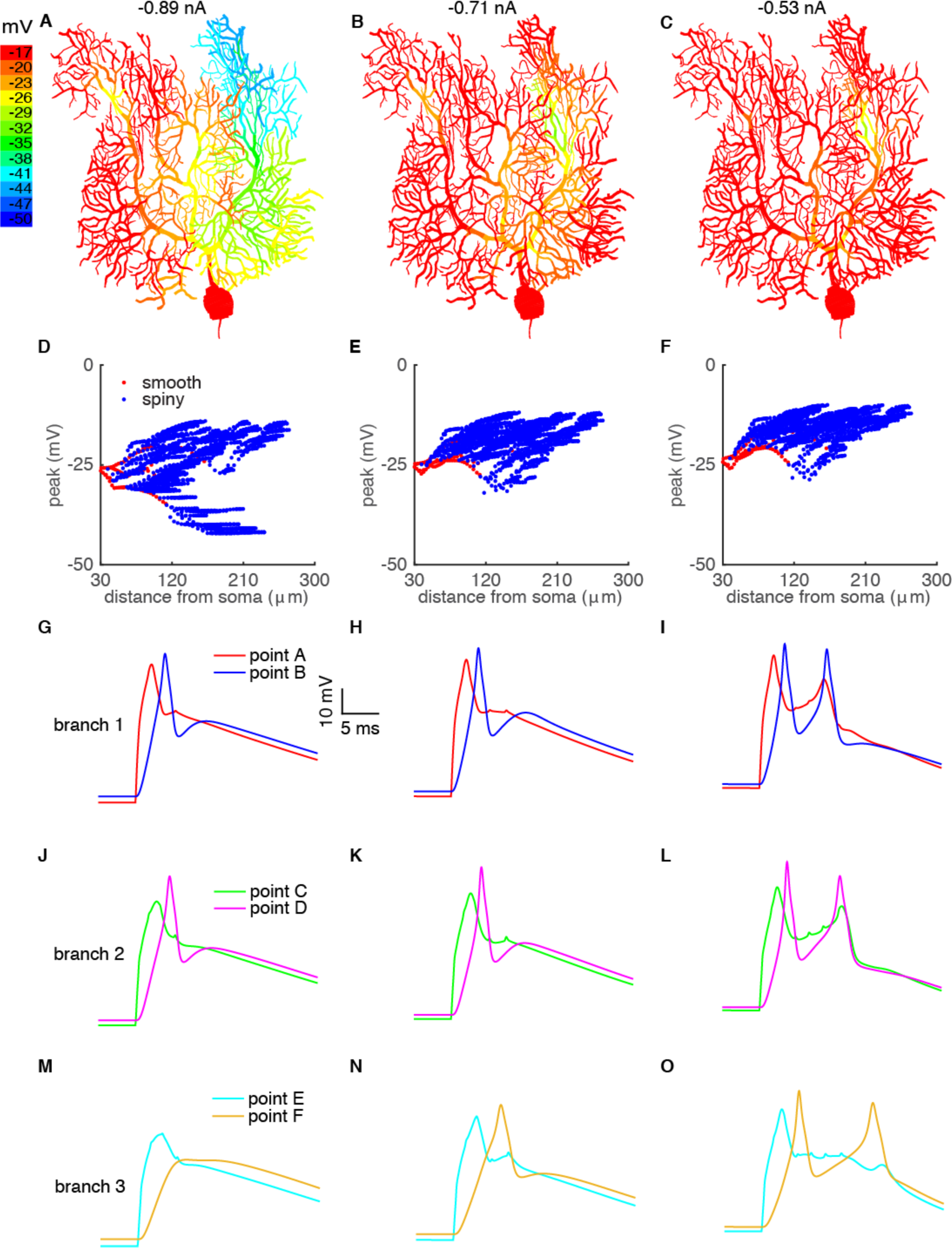
Locking Kv4 current facilitates dendritic spike generation and propagation and makes dendritic excitability more homogeneous. Related to Figure 3 and Figure 4. Dendritic responses to holding currents of -0.89 nA, -0.71 nA and -0.53 nA are shown from left to right. Color-coded voltage response, distance-dependent propagation of voltage response, and voltage responses (EPSP or spike) at different branches are shown from top to bottom. The definition of recorded dendritic sites is the same as in Fig. 2.

**Fig. S4.**
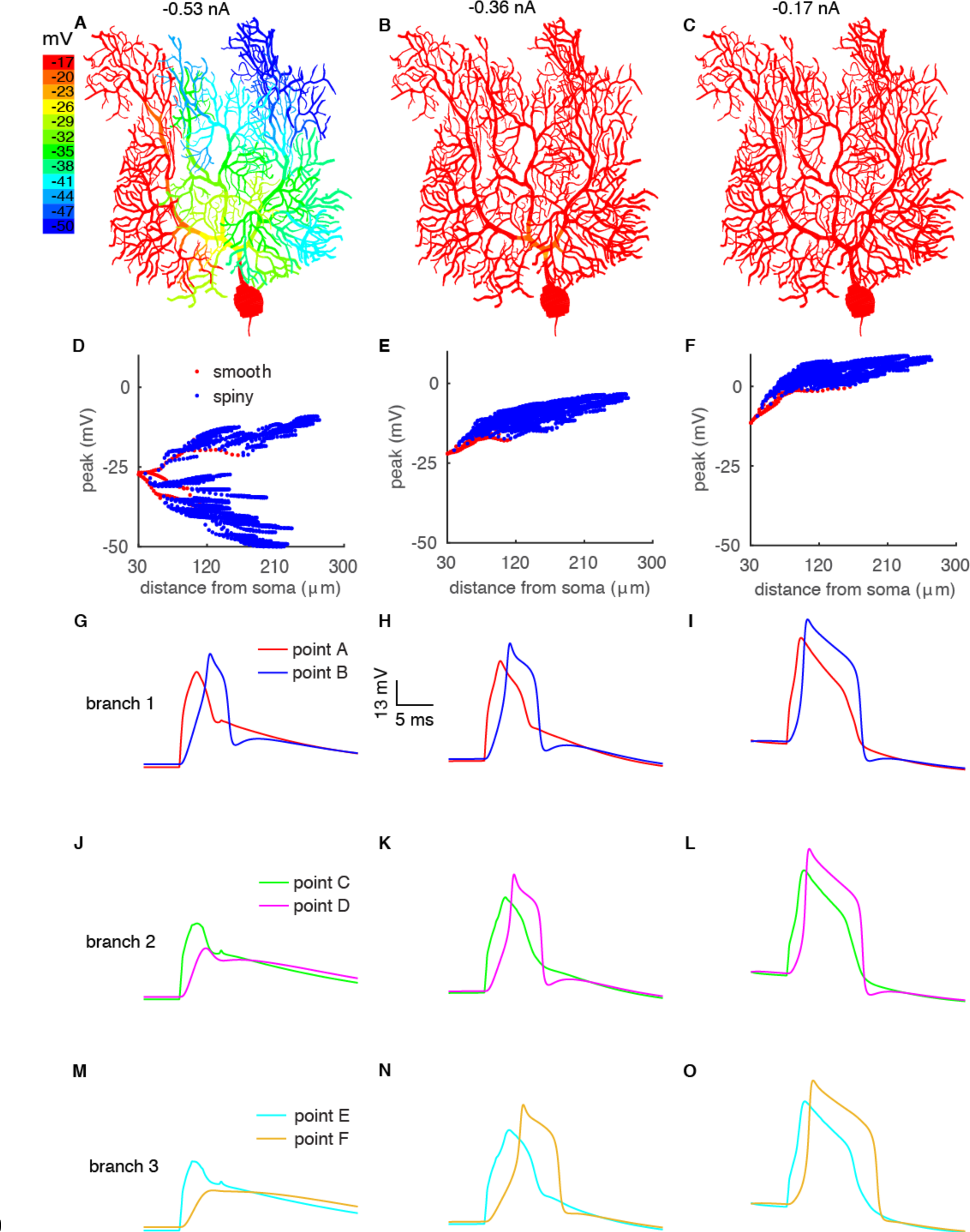
Blocking Kv3 current facilitates the propagation of dendritic spikes by increasing their durations. Related to Figure 3. Dendritic responses with holding currents of -0.53 nA, -0.36 nA and -0.17 nA are shown from left to right. Color-coded voltage response, distance-dependent propagation of voltage response, and voltage responses (EPSP or spike) at different branches are shown from top to bottom. Notice the broader dendritic spikes (G-O) with an overshoot amplitude (dendritic peak is above 0 mV in F). The definition of recorded dendritic sites is the same as in Fig. 2.

**Fig. S5.**
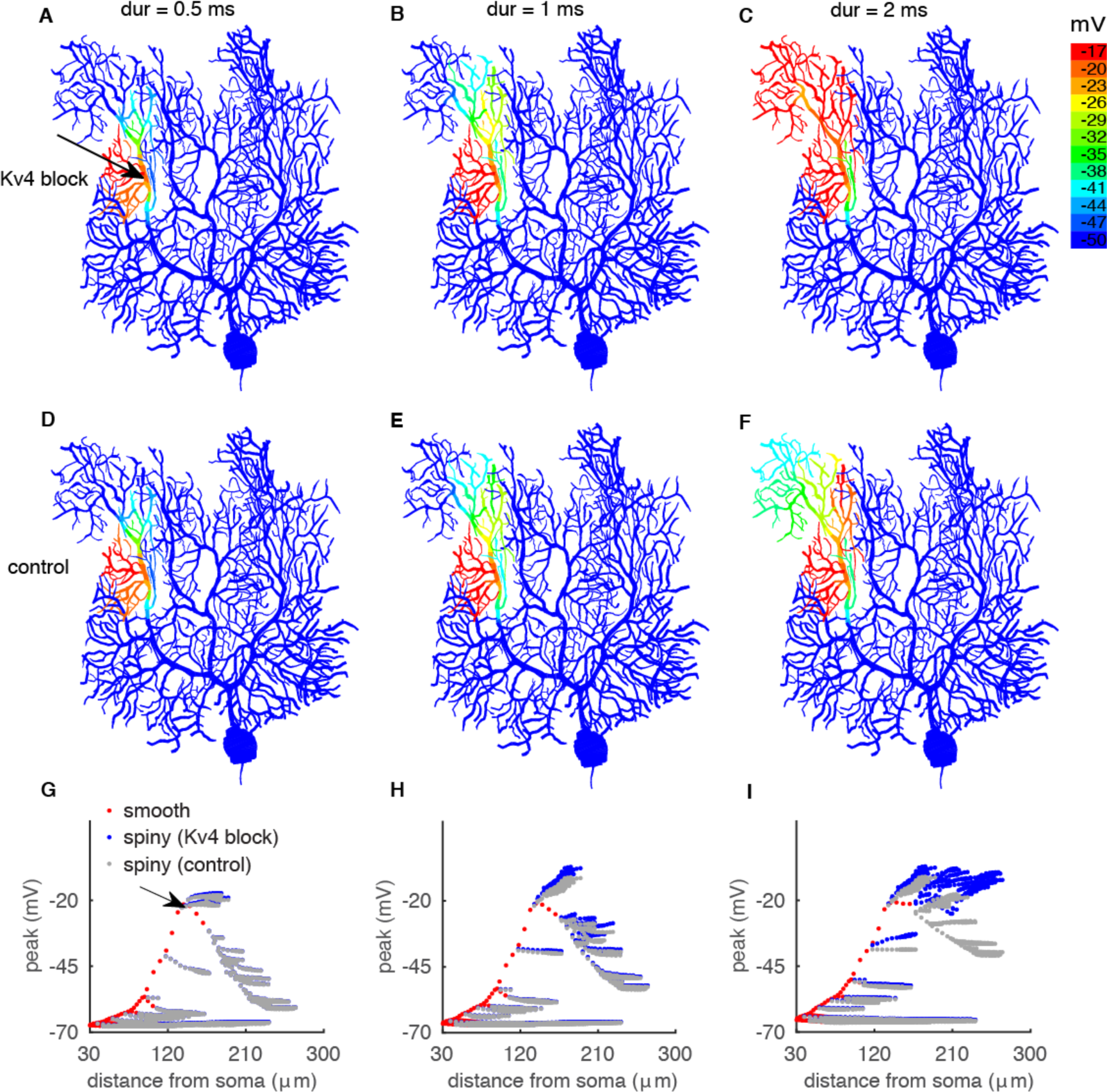
Broadening of dendritic spikes facilitates their propagation. Related to Figure 3. We exerted square wave voltage clamps (peak is -20 mV, similar with dendritic spikes) with varying durations at the indicated sites of the smooth dendrite to explore propagation efficiency. The duration increases from 0.5 ms to 2 ms from A to C (Kv4 block) and D to F (control). The distance-dependent propagation of this signal is shown in G-I to reflect the relative contribution of impedance mismatch and Kv4 current on the decay of this signal. The clamped site is indicated in G. The PC is voltage clamped to -70 mV at the soma. Notice that Kv4 block did not affect the forward propagation to the soma of this signal.

**Figure S6.**
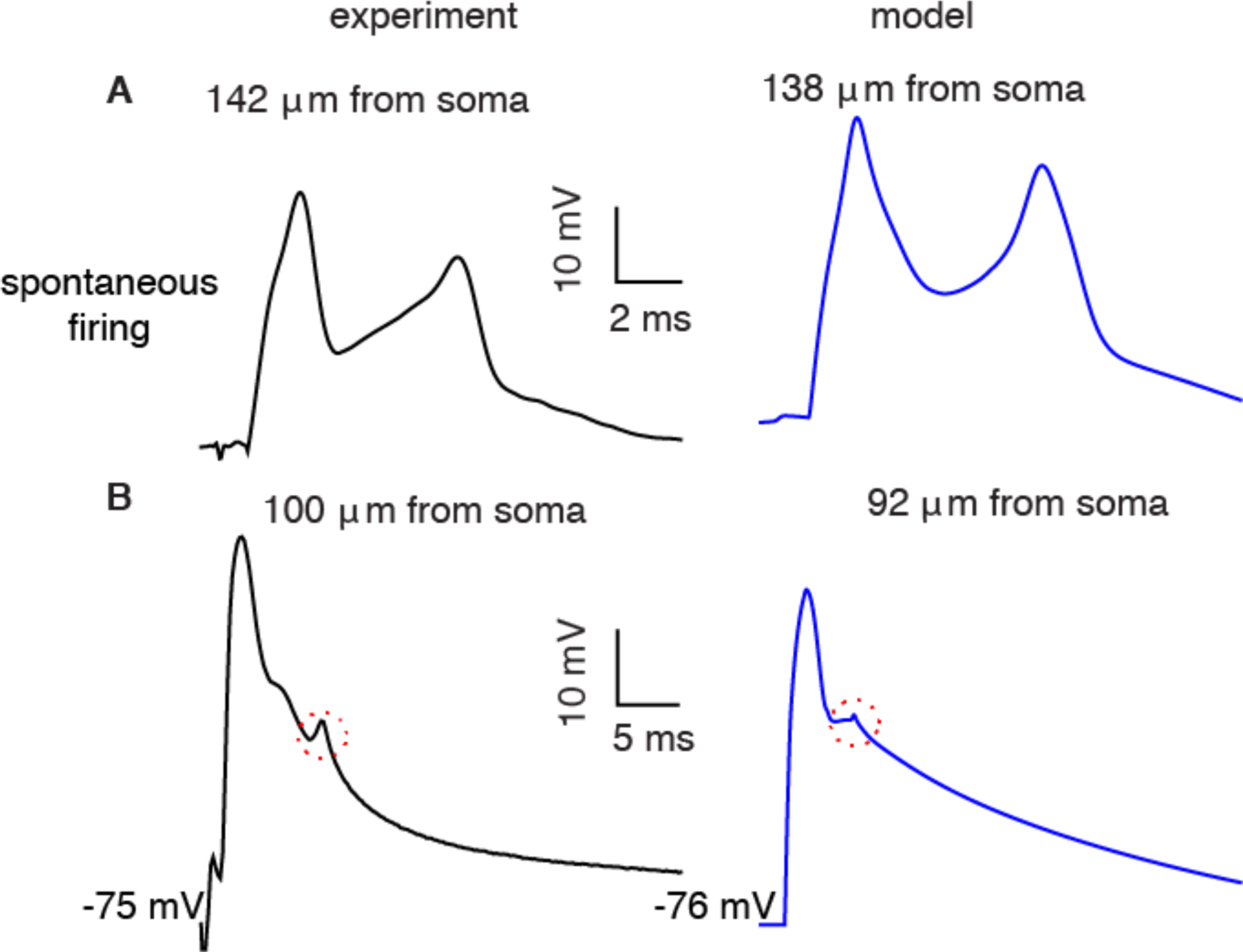
CF-evoked dendritic voltage response waveforms with different current injections (holding potentials). Related to Figure 3. A. CF-evoked dendritic spikes with two spikelets in the experiment (left, reproduced from Fig. 5A of Davie et al. (2008)) and in the model (right). B. CF-evoked dendritic voltage responses in the experiment (left, data from Ohtsuki et al. (2012)) and in the model (right). The ‘deflection’ caused by a somatic Na^+^ spikelet can be found on top of the dendritic response under this condition in both experimental recordings and in the model (circled in the figure).

**Figure S7.**
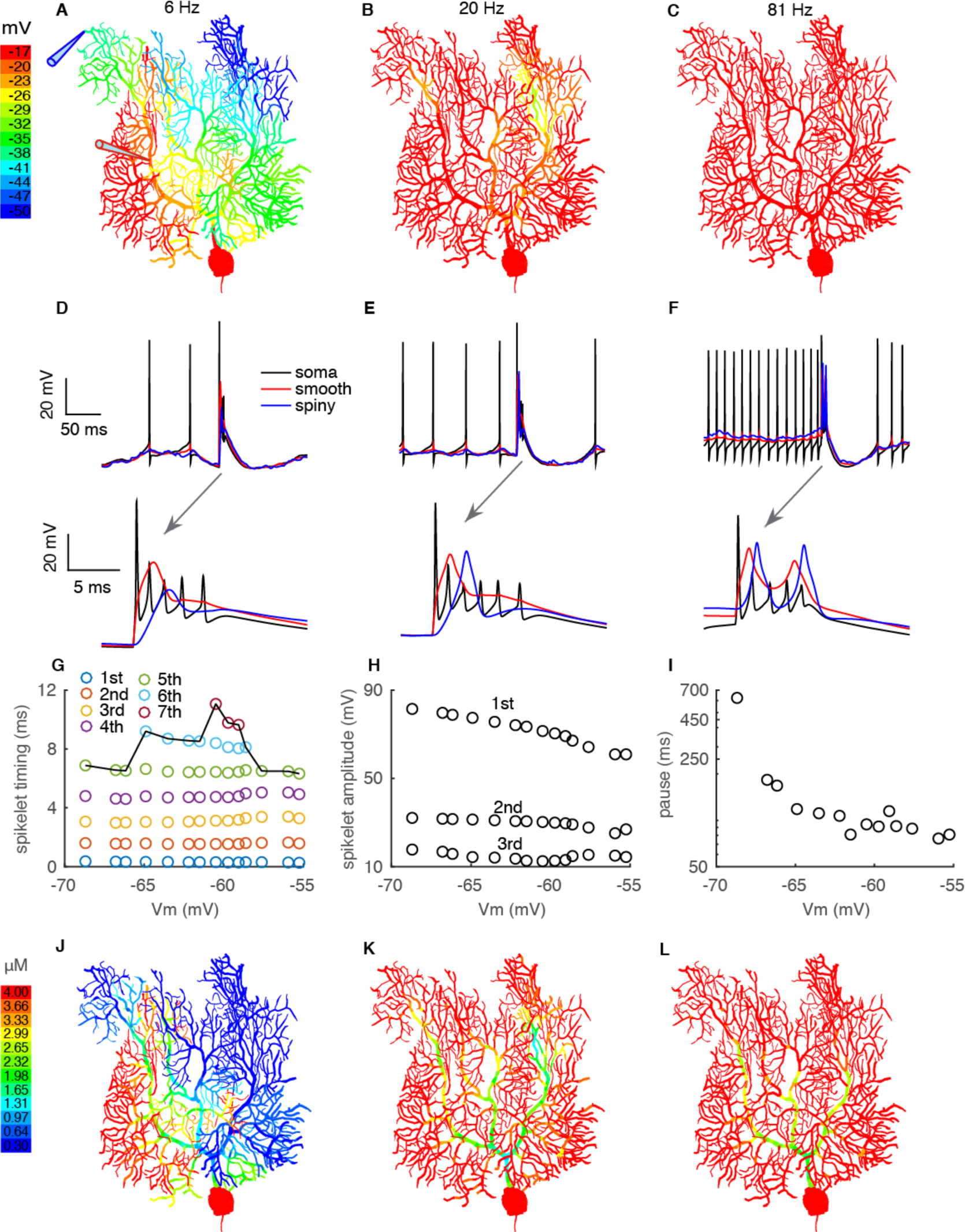
CF responses when the voltage states of the PC are manipulated by combined excitatory and inhibitory synaptic inputs. Related to Figure 5. A-C show color-coded voltage responses with increasing SSFRs. D-F show the corresponding somatic and dendritic responses. CF responses at a higher time scale are shown in their bottom panels. G shows the effect of somatic voltage states on the CS spikelet numbers and durations. H. Effect of somatic voltage states on the amplitudes of the first 3 somatic spikelets. I. CF-evoked pause decreases with depolarization. J-L show color-coded Ca^2+^ concentrations with increasing SSFRs.

**Figure S8.**
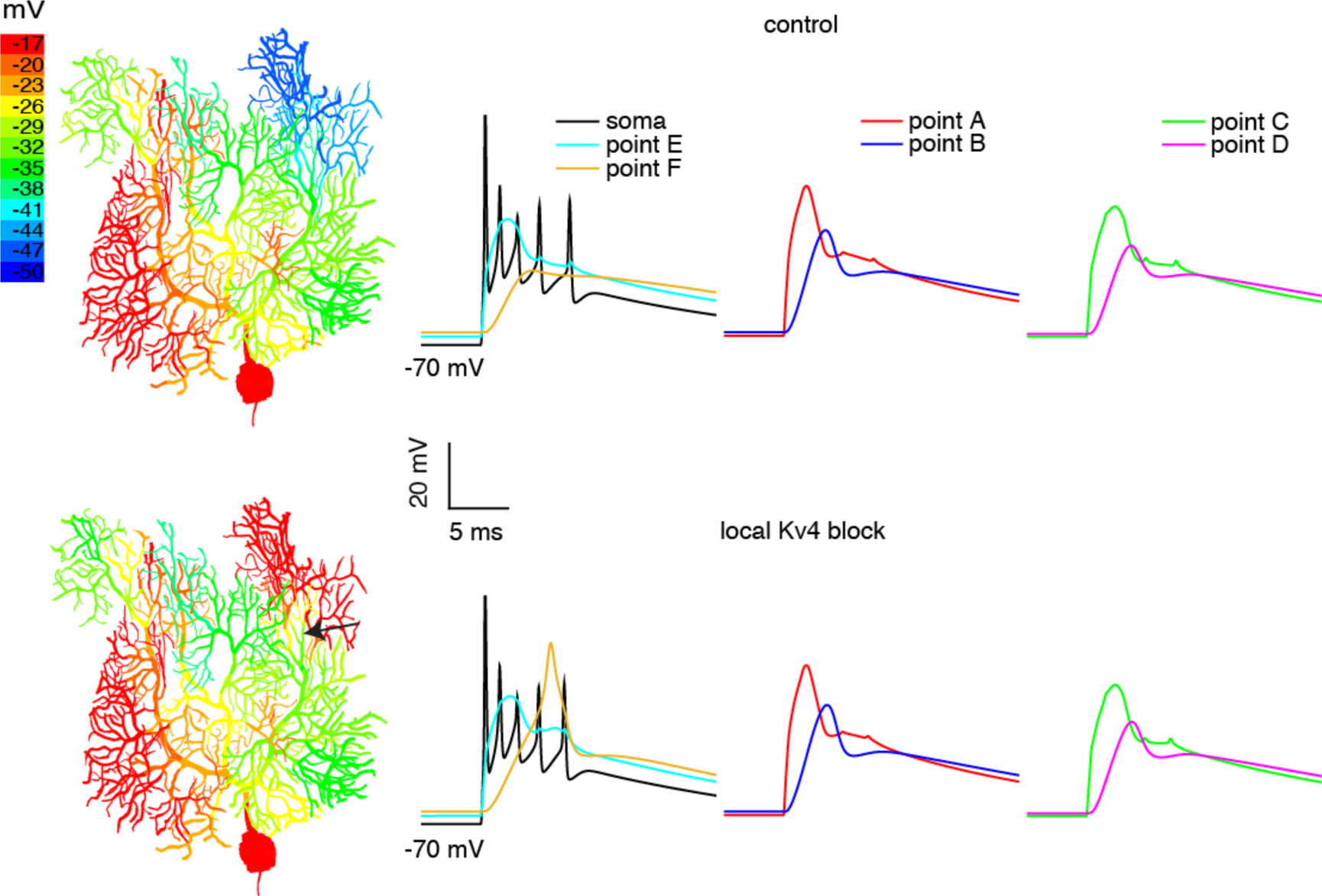
Local Kv4 block modulates dendritic responses in a spatially precise manner. Related to Figure 6. In the child branchlet of the labeled dendritic section, Kv4 current is blocked (bottom panel). Only the dendritic responses located within the labeled branchlet (For example, Point F) are facilitated by the local Kv4 block. In other parts of the dendrite, dendritic responses are not affected. The definition of recorded dendritic sites is the same as in Fig. 2.

**Figure S9.**
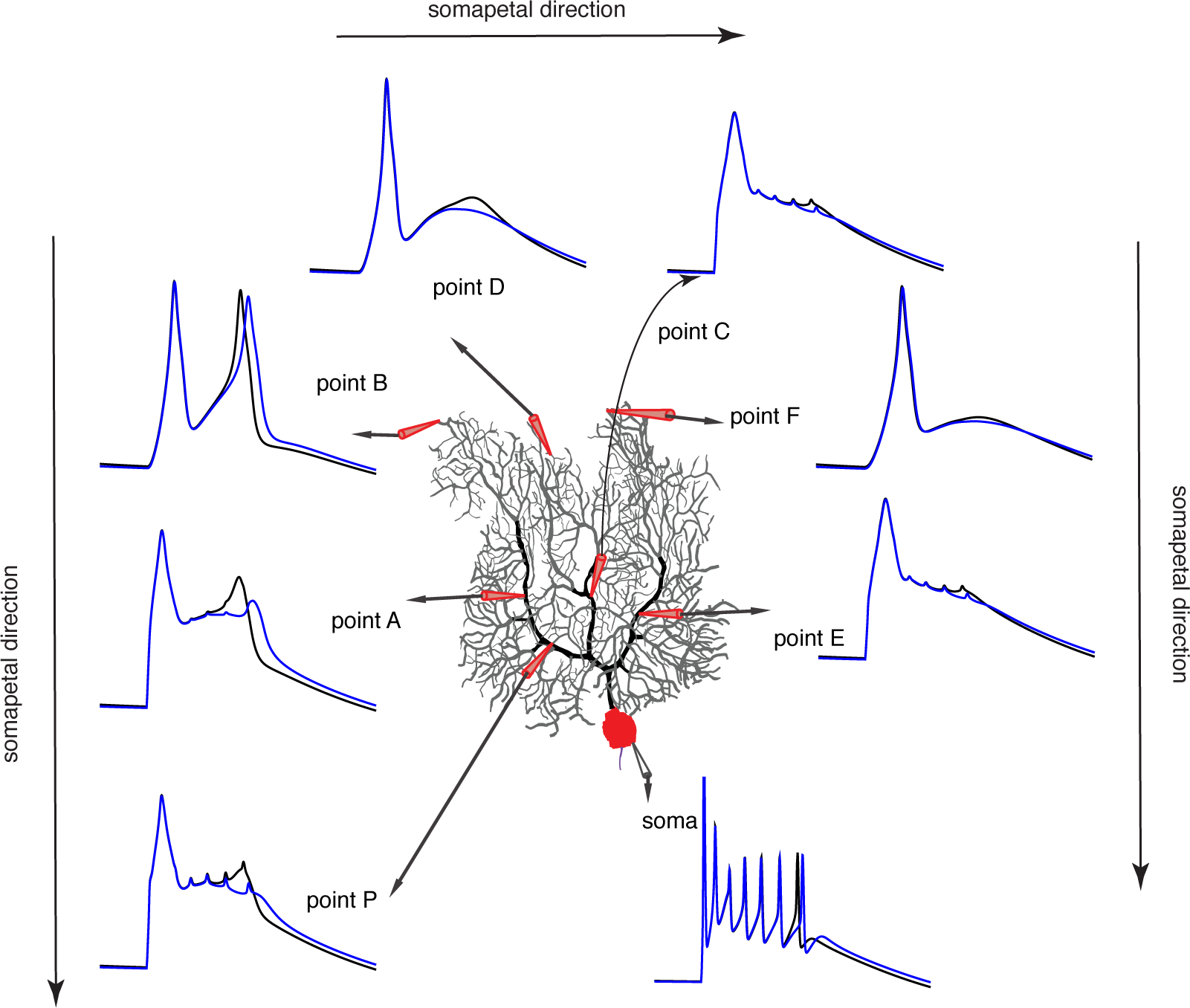
The dendritic spike measured on the proximal smooth dendrite can’t reflect dendritic responses in the rest of the dendrite. This figure supports our results in Fig. 5 and Discussion. Here we reproduce the experiment by Davie et al. (2008). The holding current is -0.2 nA. The CF is activated at different phases of the somatic interspike interval to achieve a slight change of the dendritic membrane potential. In case I (blue traces), a secondary dendritic spikelet occurs at point B of the distal dendrite. However, at point A and P of the same branch (branch 1) closer to the soma, no secondary dendritic spikelets are triggered. With a slight depolarization of the dendrite in case II (black traces), secondary dendritic spikelets occur at both proximal sites (point A and point P) and at the distal site (point B).

## Supplementary Experimental Procedures

### Current equations and calcium handling

#### 1) Na^+^ current in the PC

Na^+^ current in the model is distributed in the AIS and soma. Na^+^ current in the PC is composed of ‘transient’, ‘persistent’ and ‘resurgent’ components. We started from the previous model (Khaliq et al., 2003) and made slight changes (Narsg). In the soma, we adjusted the reaction rates between the I1 - I5 states by right shifting 5 mV (Yan et al., 2014): *α =* 150 e^(v-5)/2^°, *β =* 3 e^(v-5)/-2^°. Other parameters are the same as the original model. This adjustment mainly right-shifts the inactivation curve of this current by ~ 2.5 mV. The inactivation curve of the modified model lies within measurements (Raman and Bean, 2001; Yan et al., 2014). In the AIS, in addition to the abovementioned adjustment, we also adjusted deactivation rates between C1 - C5 to lower the spike threshold in the AIS, as done by Colbert and Pan (2002). Therefore, *β = 3* _e_(^v^+i5)/-i5 for reaction rates between C1 - C5. First, by this adjustment, the contribution of AIS spikes to somatic spikes is closer to measured data (represented by *d^2^V/dt^2^* in Fig. 1D). Second, we removed the frequently failed recovery of simple spikes after a complex spike when there is no obvious after-hyperpolarization following the complex spike (although this can be observed in some experiments, data shared by Paul Mathews from UCLA).

It is impossible to approximate Na^+^ currents using a single model, given their multiple components. In the original model (Khaliq et al., 2003), the persistent component is tiny. Therefore, we incorporated an additional persistent Na^+^ current model to compensate (NaP). The steady state activation curve is from Carter et al. (2012), *m*_*ss*_ = 1/(1 + e^−(v+66)/5^). The time constant is from Baker (2005), *τ_m_ = 1/(α_m_ + β_m_)*, *α*_*m*_ = 1/(17.235/(1+ *e*^(v+27.58)/-11.47^)), *β*_*m*_ = 1/(17.235/(1+ *e*^(v+86.2)/19.8^)). This persistent component helps to maintain spike firing during negative current injections.

#### 2) H-current in the PC

H-current (Ih) is distributed in the whole PC (Angelo et al., 2007). In our model, the formulations are the same as the model formulations from Angelo et al. (2007), except we shifted the activation curve to a more hyperpolarized direction by 6 mV to make ‘sag’ responses with different holding currents more comparable to experimental recordings (Fig. 1G).

#### T-type Ca^2+^ current in the PC

T-type Ca^2+^ current is distributed in the AIS (Bender and Trussell, 2009), the smooth dendrite and the spiny dendrite (Hildebrand et al., 2009; Otsu et al., 2014). The formulation of this current is according to data from Otsu et al. (2014). *m*_*ss*_ = 1/(1 +*e*^−(v-35.606)/4.7^), τ_*m*_ = 1/(1.2757 + α_*m*_+*β*_*m*_), *α*_*m*_ = - 2.3199/(1 + *e*^(v+41.448)/-30.655)^, *β*_*m*_ = 2.5712/(1+ *e*^(v+21.7866)/-9.6306)^, *h*_*ss*_ = 1/(1 +e^(v+81.718)/6.4635^), τ_h_ = 1/(0.0076 + *α*_*h*_ + *β*_*h*_), *α*_*h*_ = 0.17746/(1 + e^(v+41.448)/-30.655)^, *β*_*h*_ = 0.13402/(1 + e^(v+94.836)/-5 5845)^. In the model, both BK and SK2 don’t sense Ca^2+^ influx from T-type Ca^2+^ channels.

#### 4) P-type Ca^2+^ current in the PC

P-type Ca^2+^ current is distributed throughout the entire PC (Indriati et al., 2013). The formulation of this current is according to the data from Benton and Raman (2009). *m*_*ss*_ = 1/(1 + *e*^−(v-25.5)/4.113)^, τ_*m*_ = 0.2 + 0.7031e^−((v+25)/14)2^.

#### 5) Ca^2+^ handling mechanism

In the dendrite, we used the formulation from Anwar et al. (2012) with 4 shells. In both soma and AIS, we made BK channels sense Ca^2+^ signals closer to the membrane surface (subspace) or Ca^2+^ channels by increasing the number of shells (Indriati et al., 2013). In the AIS, we still used the formulation as in Anwar et al. (2012), but we increased the number of shells to 10. In the soma, we used the fixed depth scheme for the outermost shell and 20 shells in total to reduce the computational load (Anwar et al., 2014). We also incorporated a sarcoplasmic endoplasmic reticulum Ca^2+^ATPase (SERCA) pump to reduce the spike-over-spike Ca^2+^ summation in AIS and soma. *J*_*SERCA*_ = 0.1 * *vol*(*i*) * *Ca*[*i*]/(*Ca*[*i*] + 0.0058(*mM*)), *vol*(*i*) means the volume of the ith shell. *Ca*[*i*] means the free Ca^2+^ concentration of the ith shell.

#### 6) BK current in the PC

We used the same formulation from Anwar et al. (2012) as the fast (or IBTX-sensitive) component of the BK current (Benton et al., 2013). We also incorporated the slow (or IBTX-insensitive) component of BK current (Benton et al., 2013) from Jaffe et al. (2011). The formulation of their type II model was used. The fast component is distributed in the whole PC and the slow component is only distributed in the soma and the AIS.

#### 7) SK2 current in the PC

SK2 current is distributed in the whole PC (Belmeguenai et al., 2010; Swensen and Bean, 2003). We used the formulation from Solinas et al. (2007) according to the data by Hirschberg et al. (1998).

#### 8) Kv3 current in the PC

KV3 current is distributed in the whole PC (Martina et al., 2003). The formulation is from Akemann et al. (2009) according to the data from Martina et al. (2007). In our model, the steady state activation curve is left shifted by 4 mV with V0.5 ≈ 2 mV, which is still within the experimental range (Martina et al., 2007; Martina et al., 2003; Zagha et al., 2010).

#### 9) Kv4 current in the PC

In our model, Kv4 current has both fast and slow inactivation components and is distributed in the smooth and the spiny dendrites (Otsu et al., 2014). We took the steady state activation curve from Gunay et al. (2008). The inactivation curve and time constants are according to experimental data of Stéphane Dieudonné. *m*_*ss*_ = 1/(1 + e^(v+49)/-12.5)^, τ_*m*_ = 1/(*α*_*m*_ +*β*_*m*_), *α*_*m*_ = 0.1342/(1 + e^(v+60)/32.19976)^, *β*_*m*_ = 0.15743/(1 + e^(v+57)/37 51346)^, *h*_*ss*_ = 1/(1 + e^(v+75.30348)/6.06329^), τ_hf_ = 1/ (α_hf_ + β_hf_), α_hf_ = 0.01342/(1 + e^(v+60)/7.86476^), β_hf_ = 0.04477/(1 + *e* ^(v+54)/-11.6315^), τ_*hs*_ = 100.

#### 10) Kv1 current in the PC

In our model, Kv1 is distributed in the dendrite (Khavandgar et al., 2005). The formulation is from Akemann and Knopfel (2006). We added an inactivation gate, according to data from Otsu et al. (2014). *h*_*ss*_ = 1/(1 + e^(v+66.16)/6.1881^). Due to the slow inactivation process, the inactivation time constant was set to be 1000 ms.

### Parameter tuning

The model was hand-tuned because extensive preceding efforts using automatic parameter searching did not achieve good results (Achard and De Schutter, 2006; Van Geit et al., 2007). Due to technical limitations, we can only get simultaneous somatic and dendritic patch-clamp recordings (single dendritic site) to constrain the model in most cases. However, it has been demonstrated that data recorded from just two locations in the same L5 pyramidal neuron are insufficient to accurately constrain a compartmental model (Keren et al., 2005). For the PC, we were able to get good F-I curves and ‘sag’ responses with automated search methods, but the dendrite model could not generate physiologic responses. This is because we don’t have sufficient data to constrain the active properties of the dendrite and the interaction between soma and dendrite by automatic parameter searching. Therefore, we decided to use hand-tuning instead, and we tried to incorporate as many experimental observations as possible into the model. After repeated modification over a period of more than two years, we achieved the present model.

#### Currents in the soma

The Na^+^ current profile nearly coincides with the total net ionic current during the depolarization phase of dissociated PC AP-clamp recordings (Carter and Bean, 2009). The total net ionic current is calculated by -C*dV/dt, where C is the cell capacitance and dV/dt is the time derivative of the voltage. The results of Carter and Bean suggest that Na^+^ current is nearly the sole current supporting the depolarization and K^+^ currents are not activated during the depolarization phase (as illustrated in Fig. 2 of their paper). This also agrees with the high activation threshold of K^+^ currents in the PC (Martina et al., 2007). This finding provided an opportunity to firmly constrain the somatic ionic currents. The measured peak Na^+^ current amplitude is 12.5 nA in dissociated mouse PCs with a capacitance of 12 pF, but the ratio of current/capacitance in the soma may be underestimated due to the possible remaining stump of the dendritic tree. In our model, we have a peak somatic Na^+^ current of 17 nA relative to a capacitance of 10.5 pF in the soma. Then we get the conductance density of somatic Kv3 current by constraining the spike half-amplitude duration (0.15 ms) close to experimental measurements (0.17 ms, we also gave consideration to the spike duration in the intact PC as shown in Fig. 1B). The high activation threshold of the Kv3 current makes it active mainly during the repolarization phase (Carter and Bean, 2009) and constrains the spike duration (Martina et al., 2007). Subsequently, we constrained the conductance density of P-type Ca^2+^ current and BK current to have an AHP (−77 mV) comparable to the recorded (−82 mV, still we also considered the AHP in the intact PC ~ -70 mV). The conductance density of P-type Ca^2+^ is constrained to be the same as that in the smooth dendrite. Then only SK2 and Ih remained unconstrained for the somatic ionic currents. We initially assigned tentative conductance density values for these two currents to develop a firing somata model (like the previous Kaliq-Raman model (Khaliq et al., 2003)). To reflect the negligible role of Ih in dissociated PC firing (Raman and Bean, 1999), the conductance density of Ih is half of that in smooth dendrite. In the intact PC, the conductance density of somatic SK2 current is adjusted to get a F-I curve close to the experimental measurements (Fig. 1G). The persistent component of Na^+^ current is constrained so that the intact PC still fires with -0.17 nA somatic holding current, but it stops firing when the current amplitude changes to -0.36 nA.

#### 2) Currents in the AIS

We constrained ionic currents in the AIS to make the model behave like experimental data in two aspects: the axosomatic delay of the simple spike (Fig. 1B) and the second derivative of the simple spike (Fig. 1D). The first reflects how the spike in the AIS precedes the somatic spike and the second reflects the contribution of the spike in the AIS to the somatic spike. We assumed the conductance densities of Ih in AIS are the same as in soma.

#### 3) Currents in the dendrite

Due to the limited quantitative data, ionic currents in the dendrite are more difficult to constrain compared with somatic currents. We first constrained the P-type Ca^2+^ current in the dendrite. In the spiny dendrite, Ca^2+^ influx during one dendritic spikelet is 1.3e-14 Cµm^3^ in our model, which is close to the experimentally estimated 1.1e-14 Cµm^3^ (Otsu et al., 2014). The CF-evoked Ca^2+^ influx increases with distance from soma when a Ca^2+^-supported dendritic spike occurs (Kitamura and Hausser, 2011; Otsu et al., 2014). Therefore, in the smooth dendrite, the current density was constrained to have a calcium influx ratio (smooth dendrite relative to the spiny dendrite at same membrane surface area) of 0.36. The conductance densities of P-type Ca^2+^ current are further supported by the time to peak of the first dendritic spikelet being ~ 1.5 ms in the smooth dendrite (Davie et al., 2008; Kitamura and Hausser, 2011). Conductance densities of Kv4 current are assumed to be homogeneously distributed and then repeatedly adjusted to make this current able to ‘brake’ the dendritic spikes at low voltage ranges. We distributed more T-type Ca^2+^ channels (Hildebrand et al., 2009) in the spiny dendrite given they are preferentially distributed on spines. This current helps to offset the window current of Kv4 current to maintain the dendritic interspike interval membrane potential. We distributed larger conductance densities of BK current on the smooth dendrite compared to on spiny dendrite, given the BK channel clusters are mainly distributed on the soma and smooth dendrite (Kaufmann et al., 2010). BK current contributes to reasonable AHP after complex spikes as found in Davie et al. (2008). Ih is constrained to get a ‘sag’ response close to the experimental recordings (Fig. 1G). Kv1 and SK2 help to regulate dendritic excitability to stop the occurrence of Na^+^-Ca^2+^ bursting at low current injections. In our model, as with most cell recordings in rat PCs, Na^+^-Ca^2+^ bursts begin to occur with 1.25 nA somatic current injection. In addition, this dendrite model can replicate the distance-dependent decay of the simple spike amplitude (Fig. 1E), the regulatory role of K^+^ currents in voltage-dependent dendritic spike generation and propagation (Fig. 3, S2-S4, S7), the dendritic voltage response waveforms at different conditions (Fig. S6), and peak dendritic voltage response propagation with distance from the soma at specific condition (Fig. 3G).

### Current conductance densities in different parts of the PC model

| Currents (nS/$cm^2$) | AIS | soma | smooth dendrite | spiny dendrite |
| --- | --- | --- | --- | --- |
| Narsg | 0.56 | 0.0317 | 0 | 0 |
| NaP | 0.0023 | $1.4 \times 10^{-4}$ | 0 | 0 |
| $I_h$ | $1.08 \times 10^{-4}$ | $1.08 \times 10^{-4}$ | $2.16 \times 10^{-4}$ | $3.24 \times 10^{-4}$ |
| T-type $Ca^{2+}$ | $1.28 \times 10^{-4}$ | 0 | $2.7 \times 10^{-5}$ | $1.08 \times 10^{-4}$ |
| P-type Ca<sup>2+</sup> | 1.05 | 1.9e-4 | 1.9e-4 | 7.6e-4 |
| BKfast | 6 | 0.8736 | 0.21 | 0.0448 |
| BKslow | 1.05 | 0.3 | 0 | 0 |
| SK2 | 0.0278 | 0.0075 | 2.16e-4 | 3.6e-4 |
| Kv3 | 115 | 1.8 | 0.1512 | 0.2268 |
| Kv4 | 0 | 0 | 0.015 | 0.015 |
| Kv1 | 0 | 0 | 0.002 | 0.001 |

